# SaVanache: indexing and visualizing pangenome variation graphs

**DOI:** 10.64898/2026.05.05.722901

**Authors:** Mourdas Mohamed, Éloi Durant, Mathieu Rouard, Cédric Muller, Cécile Monat, Matthieu Conte, François Sabot, the SaVanache Consortium

## Abstract

With the rapid increase in genome sequencing and the growing availability of genomic resources, genomics is shifting toward pangenome representations that capture intra- and inter-specific diversity by integrating multiple genomes into a single entity. These pangenomes are increasingly modeled as graphs, encoding complex genomic variations in structures such as de Bruijn or variation graphs. However, while genome browsers provide standard and effective solutions for visualizing single or limited numbers of genomes, equivalent interactive tools for graph-based pangenomes remain limited, particularly for variation graph models.

We developed SaVanache, a multi-resolution visualization interface designed to explore pangenome variation graphs at various depths. SaVanache enables the exploration of both global diversity and structural variations (SVs) across genomes relative to a user-defined linear pivot genome. Unlike synteny viewers, SaVanache emphasizes variations by representing SV types through a dedicated set of glyphs, facilitating intuitive one-to-many comparisons. To support smooth exploration, SaVanache preprocesses a Graphical Fragment Assembly (GFA) pangenome file into optimized index and data structures, enabling fast, real-time queries on large pangenome graphs.

By combining advanced visualization techniques with efficient data handling, SaVanache provides a robust tool for scientists to analyze and visualize genetic variation within genomes and pangenomes.

SaVanache is open source and freely available at https://forge.ird.fr/diade/savanache, with a public web instance at https://savanache.ird.fr.

**Author summary:** Pangenome variation graphs (PVGs) capture genomic diversity by integrating structural variants (SV) and single-nucleotide polymorphisms (SNPs) across multiple genomes within sequence graphs. We introduce SaVanache, a visual analysis tool designed to explore these pangenomic resources, with a focus on structural variations.

Unlike traditional genome browsers limited to a few genomes, SaVanache offers a multi-scale interface that allows users to move from pangenome diversity overviews and chromosome-scale patterns to individual structural variations and alignments. Using a user-selected linear pivot genome as an anchoring reference, SaVanache enables one-to-many comparisons against other genomes, simplifying complex PVG structures into comparable units. Structural differences are represented using a visual representation based on glyphs to distinguish different types of rearrangements between otherwise syntenic genome segments. Built with real-time data updates in mind, it efficiently handles large datasets thanks to its internal parsing and indexing backend, speeding up data retrieval.

SaVanache supports interactive diversity scatterplots, dynamic switching of the pivot genome, grouping genomes by metadata to explore genotype-phenotype links, navigate through annotation, and a local graph pivot-based vizualization. Complete with export functions, it bridges visual exploration to downstream bioinformatics analyses. By combining these complementary views, we aim to make structural variations in PVGs easier to explore and compare across many genomes, bringing new ways of exploring and interacting with pangenomes.

## Introduction

Over the last 15 years, genomics has undergone a major transformation driven by the successive appearance of second then third generation of sequencing technologies, together with their increasing affordability. This led to an explosion in the number of sequenced genomes, first in organisms characterized by relatively small, structurally simple, and compact genomes such as bacteria, and later in larger and more structurally complex eukaryotic genomes [1, 2]. In addition, multiple individuals per species were increasingly sequenced in order to characterize the fullest genetic and genomic diversity contained within each species or group. For many years, this diversity was mainly characterized through single nucleotide polymorphisms (SNPs) due to the massive use of short read sequencing. However, access to long-read technologies now allows more reliable detection of large structural variations (SVs) across many genomes. Although less numerous than SNPs, SVs can range from a few nucleotides to hundreds of kilobases and have been proven to have impact on numerous phenotypes [3, 4].

Genomic variation was initially characterized through comparison of sequencing reads to a single individual identified as a well-assembled ‘reference genome’. The increasing number of whole-genome assemblies available within the same species has revealed the limitations of this approach, particularly in capturing structural differences among individuals [5–8]. The study of this diversity gave rise to the concept of ‘pangenome’, which aims to represent the full range of genomic diversity within a species or a defined group [9]. Initial studies focused mainly on pangene sets, studying gene content variations *via* copy number or presence/absence variations. More recently, pangenomics has expanded towards genome-wide variations, combining SNP and SV into structured representations known as pangenome graphs. Earlier graph representations, still widely used, often relied on *k*-mers (like de Bruijn graphs), while pangenome variation graphs (PVGs) have become increasingly used for large eukaryotic genomes [10]. PVGs are generally directed graphs representing compacted whole-genome alignments where nodes are sequences present in at least one haplotype and edges represent adjacencies observed in one or more haplotypes. Individual genomes and haplotypes can then be reconstructed as paths (or walks) through the graph.

As they derive from whole genome alignment, PVGs incorporate both SNPs and SVs in their structure [11–13]. However, extracting and interpreting these variations directly from the graph remains a challenge, even when the graph is encoded in a GFA file (Graphical Fragment Assembly, a standard format used to store genome graph information). For biological interpretation, these variations need to be identified, compared and explored efficiently. In addition, while PVG models are exhaustive data structures convenient for storage and manipulation, their visualization is complex in their native graph form (leading to the so-called “Hairball effect”),especially when many genomes, alternative paths and rearrangements are simultaneously represented.

As pangenomes became more widespread, visualization tools evolved from gene- and alignment-centered approaches such as Pan-Tetris [14], PanViz [15], panX [16], and PanVA [17], to embrace representations adapted to pangenome graphs. Graph viewers such as Bandage [18] and GfaViz [19], as well as dedicated tools such as ODGI [20], SequenceTubeMap [21], and PanGraphViewer [22], provide direct access to graph topology and genome paths. However, displaying many nodes, paths, and rearrangements simultaneously can rapidly reduce visual readability.

Alternative approaches avoid exposing graph topology directly and instead rely on linear, anchored, or browser-like abstractions. Panache [23], PPanG [24], Pantograph [25], VRPG [26], Panagram [27], and PangyPlot [28] provide different ways to summarize or navigate pangenomic variation across scales. Tools more specifically dedicated to structural variations also exist, such as MoMI-G [29] for the exploration of complex variants, or plotsr [30] for the visualization of syntenic rearrangements. A comparison of these tools, their visualization paradigms and input requirements is provided in the Supplementary file **??**.

Although the landscape of tools for visualizing structural variations is diverse [31], existing approaches generally emphasize sequence similarity, synteny, conservation or local structural changes, with limited characterization of variations. Comparing structural differences across many genomes while preserving their genomic context and conveying biological relevance remains challenging.

Derived from Panache [23] which was focused on pangene sets, SaVanache is a visualization framework for pangenome variation graphs dedicated to the exploration of structural differences between genomes. It introduces (i) a multi-scale exploration workflow spanning from pangenome diversity overviews to chromosome-level structural variation inspection, (ii) a dynamic pivot-based comparison enabling flexible one-to-many genome exploration, and (iii) a novel glyph-based visual grammar for structural variation designed to facilitate the interpretation of complex structural differences embedded in the PVG. SaVanache further supports access to the corresponding graph paths and sequence extraction, the navigation through a provided gene annotation, and relies on efficient indexing to enable responsive interaction with large pangenome datasets.

SaVanache is written in JavaScript and Python. It is part of the tools from the GraSuite (https://forge.ird.fr/diade/grasuite), licensed through GNU GPLv3 and available on the IRD Forge (https://forge.ird.fr/diade/savanache). A Docker file allows an easy deployment, and a public instance is available with various PVGs examples at https://savanache.ird.fr.

## Results

### SaVanache: multi-level visual exploration

SaVanache provides a set of multiple complementary views designed to support the exploration of PVGs at multiple scales. Recognizing the wide range of information required to explore the diversity of one or several pangenomes, we designed SaVanache as a multi-page viewer. This multi-page browser allows progressive zoom-in and zoom-out during PVG exploration, avoiding to overwhelm users with a unique page containing excessive information. This design enables users to progress through their exploration *via* interconnected pages, diving deeper into genetic analysis while maintaining the ability to navigate backwards to modify their dataset selection. The successive pages first allow users to explore the available PVG catalog, then to examine and to compare their genetic diversity, and finally to investigate from single base to chromosome-level structural variations. These different view levels, available in a single tool, offer users the freedom to navigate complex genomic datasets, to examine variation patterns, and to export sequences or alignments for downstream analyses.

### Time and memory

SaVanache relies on the indexing of the GFA 1.1 in different subfiles (see Design & Implementation section below), and thus requires CPU time, Memory usage and disk space. As an indicative benchmark, we tested different configurations on the same machine (Intel Xeon Gold 2.20 GHz 6238R CPU, 112 threads, 64 Gb RAM):

- *Escherechia coli*, 13 samples, 30 cores, 55sec, 345Mb RAM, 113Mb disk space (including PVG)
- *A. thaliana*, 69 samples, 10 cores, 2h 41mn 25sec, 5Gb RAM, 122Gb disk space (including PVG)
- *A. thaliana*, 69 samples, 30 cores, 2h 06mn, 5Gb RAM, 122 Gb disk space (including PVG)
- *Oryza sativa*, 13 samples, 30 cores, 36mn 23sec, 3.7Gb RAM, 154Gb disk space (including PVG)
- *Homo sapiens*, 10 samples/Chromosomes 1 to 7, 50 cores, 6mn, 20 Gb RAM, 5.8Gb disk space (excluding PVG)
- *H. sapiens*, 45 samples, all chromosomes, 30 cores, 6h 30mn, 40Gb RAM, 1Tb disk space (including PVG)

However, these time and memory consumptions depend more of the node depth and number rather than the graph size itself.

### A catalog of PVGs

The first view of SaVanache displays the full list of PVGs (implemented in GFA1.1 format) available in the system, most of which were generated as part of this work, offering users a curated and readily explorable collection of pangenomes across species using the online version (Figure 1). It enables users to identify all available pangenomes for a given species or group of genotypes, and provides a quick description of each one. Indeed, as the number of pangenomes increases with different sampling and sources, it is very likely that users would like to explore different pangenomes from the same species, depending on their analyses and research projects. Then, by selecting one or several pangenomes, users can proceed to the next page which allows them to describe in more detail the genomes present in each pangenome. This first view also provides a synthetic view of the PVG size depending on metadata (e.g. name, classification).

**Fig 1.**
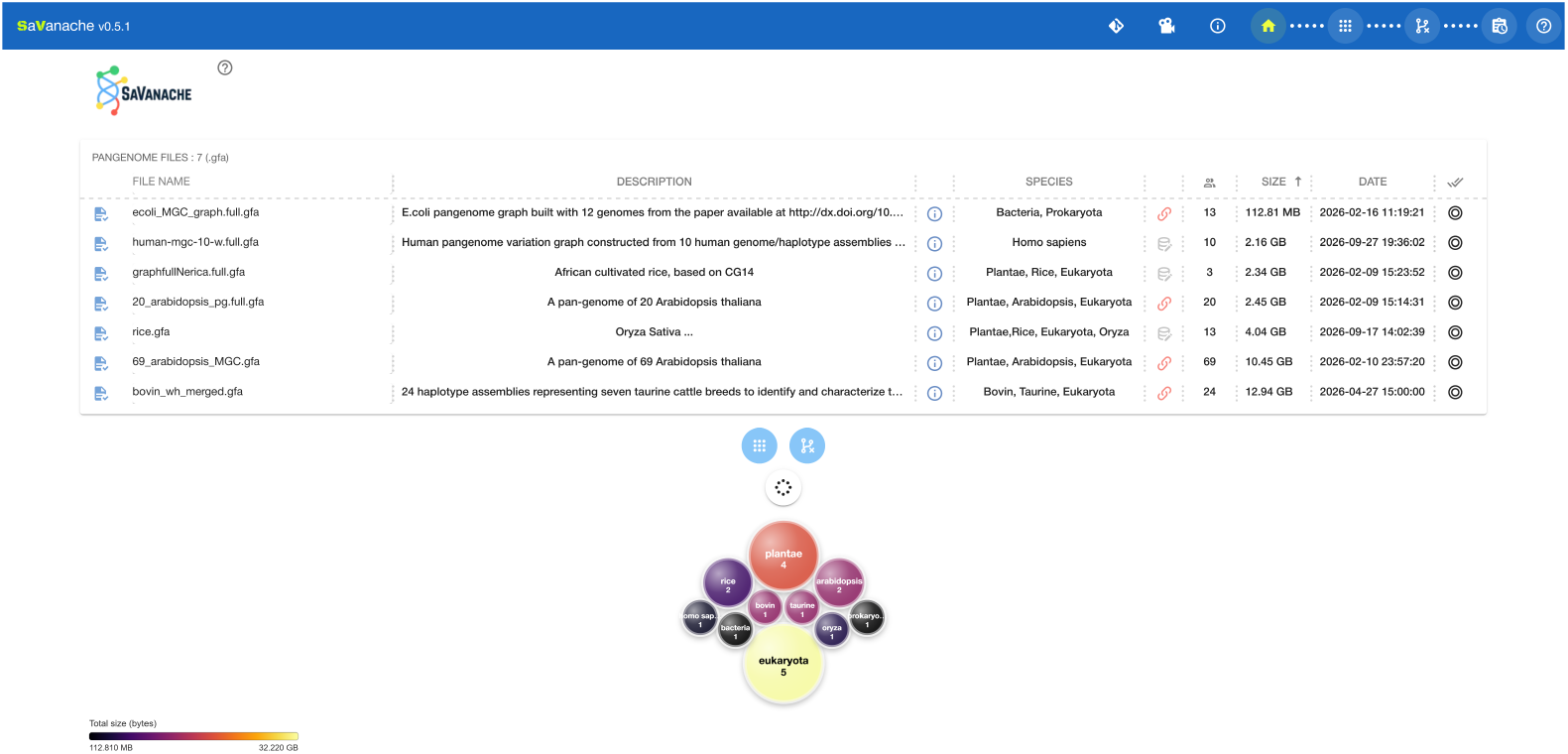
Overview of pangenome variation graphs (PVGs) available in the public SaVanache implementation. The upper panel lists indexed PVGs with associated metadata (description, species, size, and date), including multiple pangenomes for the same species (e.g., *Arabidopsis thaliana* or Asian rice), enabling comparison of different sampling strategies and pangenome constructions. The lower panel summarizes dataset composition using a node-based visualization, where each node corresponds to a metadata category and node size reflects the cumulative size of the associated PVGs.

### Diversity overview in PVGs

The second view of SaVanache displays an interactive scatterplot of all available genomes in the selected PVGs (Figure 2). Individual genomes are represented as either dots or diamonds (the latter when they were used as the initial seed to create the PVG, or indicated as ‘reference’ in the GFA file), positioned according to the first two dimensions of a MDS analysis (see Methods). The more similar the genomes are, the closer they are in the plot. Genomes are additionally colored based on the PVG (standard color panel) they belong in the dataset. When hovering over a genome circle (or diamond), all other genomes present in the same PVG are highlighted, enabling further inspection of intersections between pangenomes (Figure 2). This Panel summarizes then the high-dimension diversity of all uploaded genomes and helps users select the most relevant PVG for their analysis.

**Fig 2.**
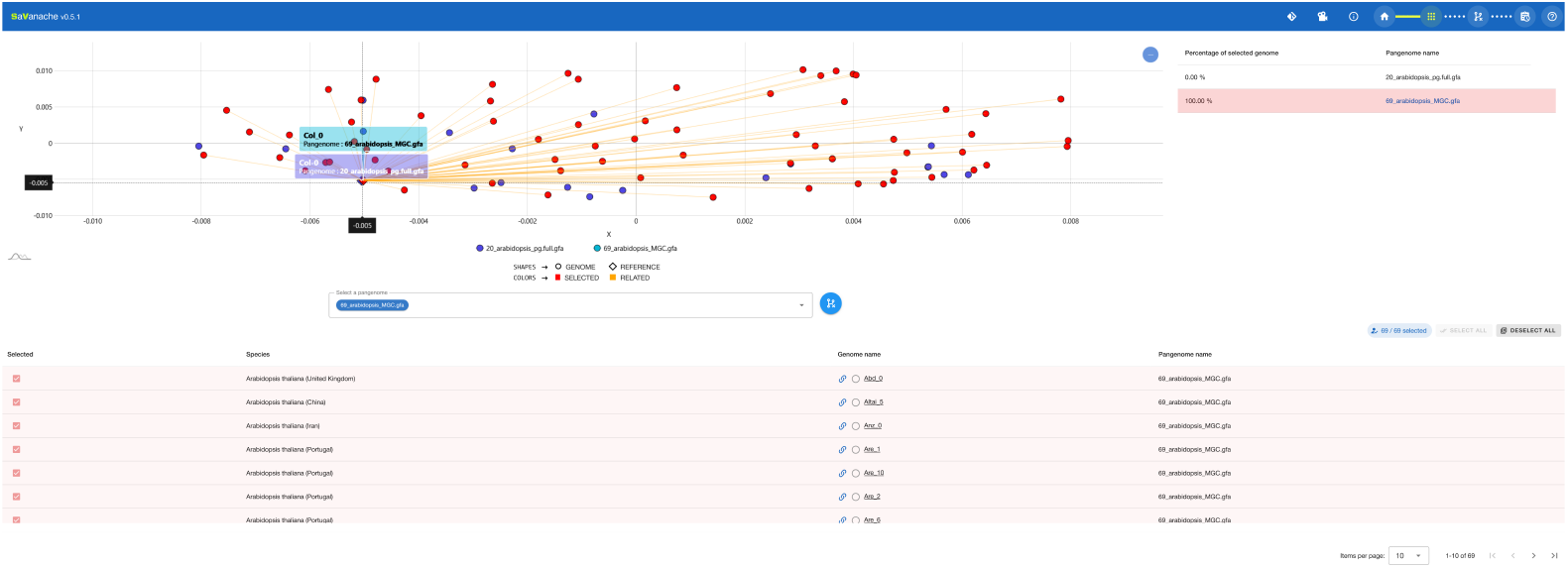
Diversity panel overview of genomes embedded in the selected PVGs. Genomes from each PVG are projected on a shared multidimensional scaling (MDS) based on their genomic similarity (*k*-mer-based), allowing direct comparison across pangenomes. The genome assembly (Col-0) is positioned near the plot origin in both the 20_arabidopsis_pg.full.gfa and 69_arabidopsis_MGC.gfa pangenomes, reflecting their close genomic proximity and consistent placement across datasets. Related genomes (orange) radiate outward from this shared reference position, revealing that the 69-genome pangenome captures a broader range of genomic diversity than the core 20-genome set alone. Interactive links are displayed when hovering over a datapoint.

Users can thus interactively explore this space to assess relationships, clusters, or outliers. Provided metadata associated with each individual, such as external identifiers, species information, and free-text descriptions, are displayed in an interactive table below the Scatterplot. From this panel, users can also filter and select the individuals to be carried forward to the Structural Variation Browser, allowing subsequent analyses to focus on a subset of genomes of interest.

### Structural variation browser

The third view is dedicated to the interactive visualization of the SV along chromosomes between the pivot and the other genomes, in a browser-like environment (see Figure 3). Upon selection of a PVG of interest, SaVanache displays it using an abstract linear representation, displaying, for each genome, its SVs relative to a ‘pivot genome’. This pivot, which can be switched for any other available genomes in the PVG at will, acts as a linear ‘reference’ coordinate system, hence simplifying the exploration of SVs in one-to-many genome comparisons. To also address this visualization challenge, this view implements a novel visual approach using variation glyphs (Figure 4), characterizing both the relative positions and types of variations. This representation transforms complex graph divergences into clear visual elements alongside a pivot path, making SVs visually prominent. Using this one-to-many approach additionally facilitates the analysis by rendering apparent conserved or divergent patterns. Similarly to classic genome browsers, this view enables complete per-chromosome exploration, with the possibility to navigate and zoom into a particular region of interest to explore its diversity. Within such a region, users can extract sequences for each genome in FASTA format and obtain their alignment (See supplementary Figure ).

**Fig 3.**
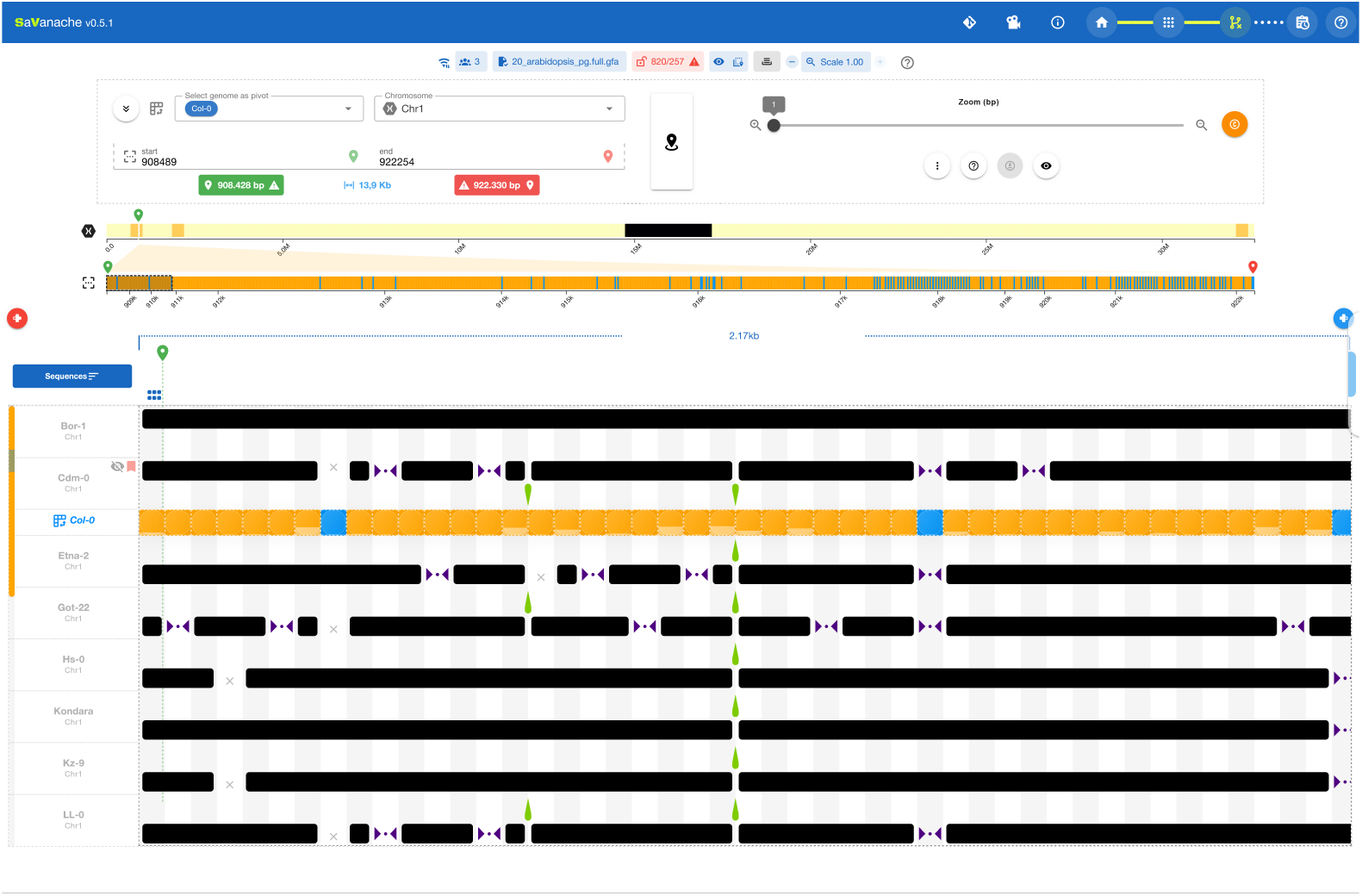
Structural Variation Browser displaying 20 *Arabidopsis thaliana* genomes. Col-0 is selected as the pivot genome. Solid black blocks indicate segments identical to the pivot, whereas dedicated glyphs represent structural variants (see text for details). The upper chromosome overview indicates the position of the displayed region within the chromosome. The current zoom threshold is set to 1 bp, corresponding to nucleotide-level resolution and allowing all graph segments of at least 1 bp to be displayed.

**Fig 4.**
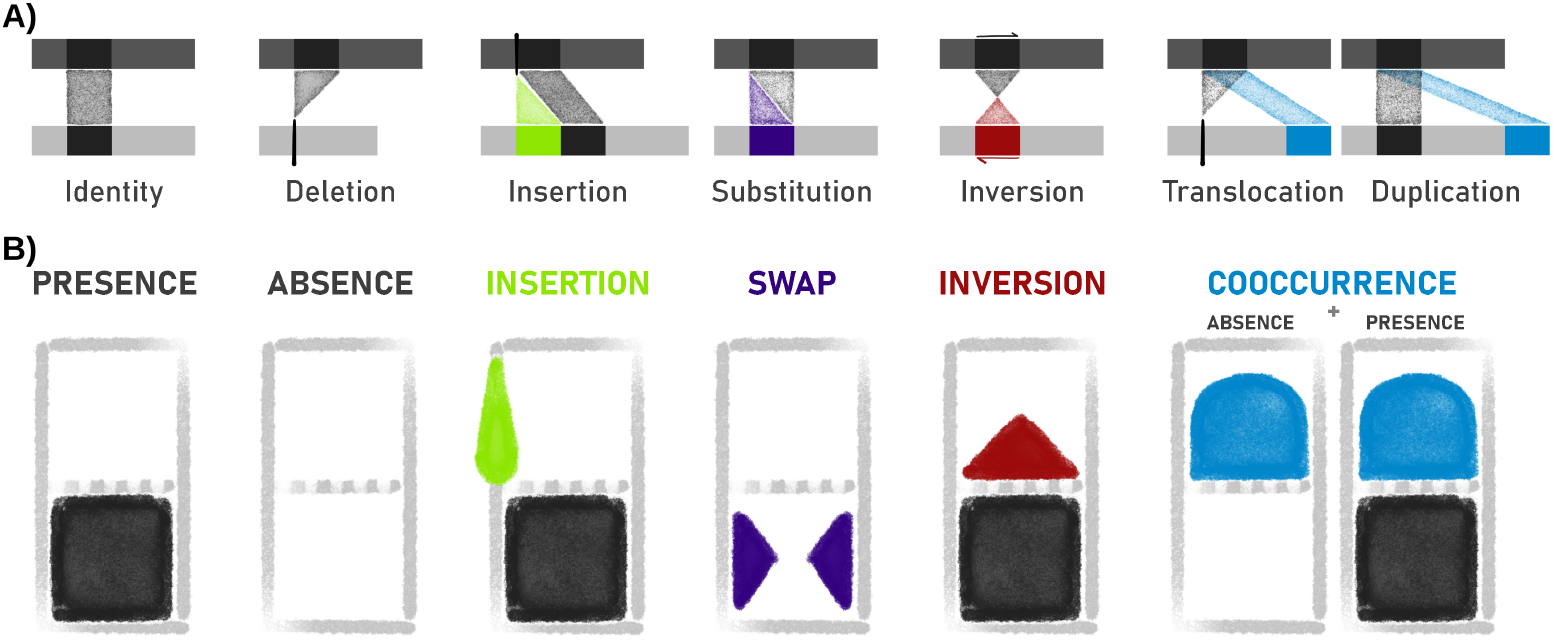
Glyph-based representation of structural variation in SaVanache. All classic structural variations can be represented in a variation glyph by combining one or more dedicated visual marks. **A)** Traditional representation of all SV types similarly to synteny viewers. The top row in dark grey depicts the pivot genome, with a segment of interest in black. Below, in light grey, is the compared genome with illustrations of each type of SV. **B)** Under each SV is depicted its equivalent using the variation glyph representation. Each visual mark encodes a type of divergence by shape, color and position (although the colors can be customized through the interface). The bottom half informs on the existence of the segment in the compared genome, the top half gives details on the succession order, direction, or possible duplicates.

### Glyph-based representation of SVs

We introduce a novel visual representation of SVs, built to highlight the differences of one genome compared to a chosen pivot. Any genomic segment (or node) of the pivot extracted from the PVG is represented as a rectangular glyph of fixed length (regardless of the true size of this node), one half informing on the presence or absence of this segment in the compared genome, the other half highlighting potential differences in synteny between consecutive segments. Within such glyphs, divergence types are redundantly encoded by specific shapes, colors, and positions, as illustrated in Figures 10 and 4. When applied to comparisons of multiple genomes to the same pivot, one per row, this compact representation enables 1-to-many comparisons and facilitates the visualization of common patterns across genomes.

Using such a representation, any usual SV can be represented in a glyph by a (combination of) visual marks encoding, in the compared Walk, as follows, the:

**◼PRES**ENCE of a segment

**◻ABS**ENCE of a segment instead, if not found at the same relative position

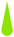**INS**ERTION of any length of genomic sequence before or after the segment

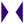**SWAP** of the segment by another sequence of any length

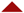**INV**ERSION of a segment

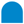**COOC**CURRENCE of a segment elsewhere

### Comparative view relative to a pivot genome

A designated *pivot genome* serves as an anchor for structural comparisons. In the lower panel of the third window (Figure 3), each row represents a genome (or haplotype), allowing users to visually inspect patterns of structural divergence across chromosomes. All SVs are extracted from the PVG and displayed relative to the pivot assembly/path, which provides a consistent baseline for comparative interpretation. Switching the pivot genome triggers an on-the-fly recalculation of all SVs, ensuring that comparisons remain accurate regardless of which individual is chosen as the ‘reference’. The pivot genome is annotated to distinguish the core components and the dispensable ones, using a customizable color coding. By default, regions present in at least 80% of genomes are classified as the core. Users may adjust this threshold, show or hide individual genomes, and group genomes by metadata categories. SV glyphs (e.g., insertions, deletions) can also be recolored through an integrated palette.

Users can further investigate individual variants by clicking on any SV marker. This action opens a dedicated view (Figure 5) that focuses on the pairwise comparison between the pivot genome and the selected genome or chromosome, enabling a detailed inspection of the corresponding rearrangements.

**Fig 5.**
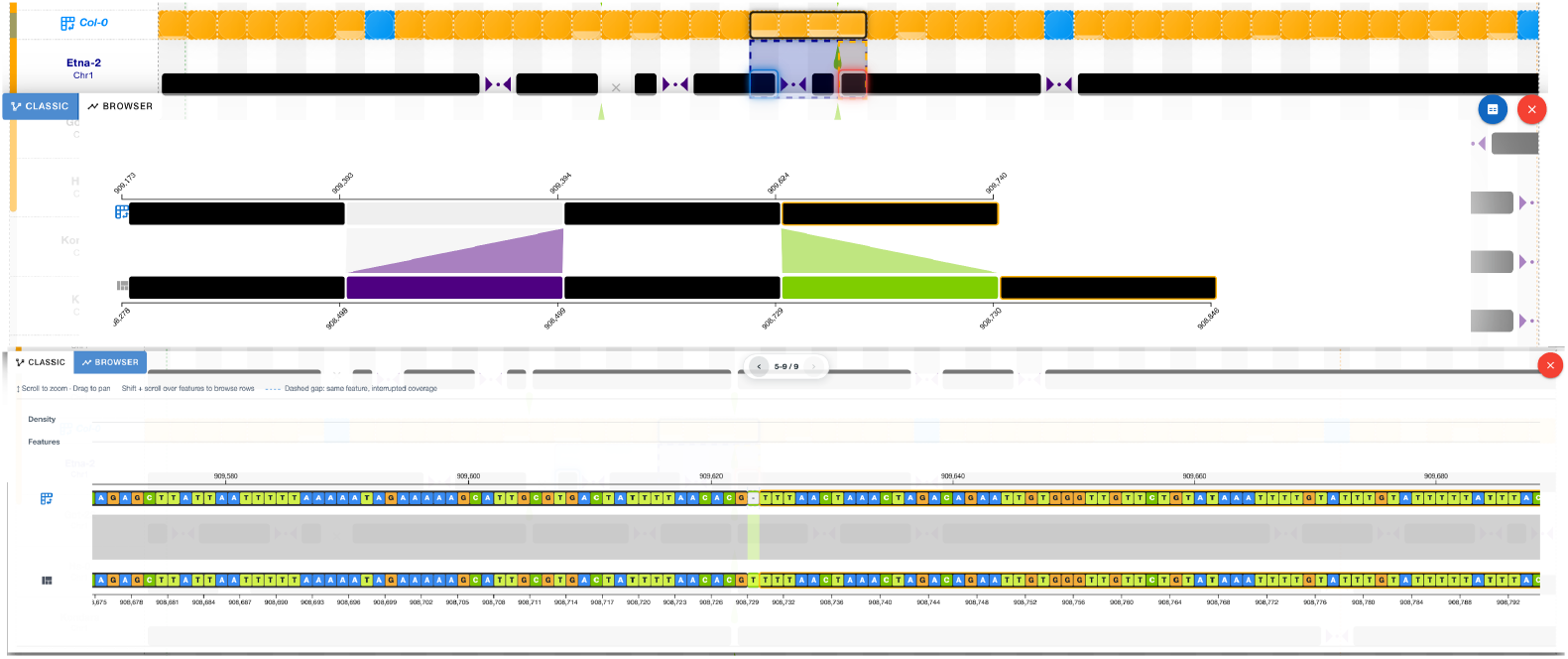
Pairwise local comparison view obtained by selecting a segment in a non-pivot genome. By clicking on any segment of a non-pivot individual, a more standard one-vs-one visualization of local variation is obtained (lower part of the figure). Here, one can see around a common region between Col-0 (pivot) and Etna-2 on the left a SNP (swap, in violet) and on the right an insertion (in green) in Etna-2. The size of the block is not correlated to the true segment size, but this can be toggled between uniform and proportional mode. For finer inspection, users can switch to a separate sequence browser, shown in the lower part of the figure. This browser displays the two sequences according to their genomic coordinates and allows continuous zooming with the mouse wheel down to individual nucleotide resolution. At this scale, SNPs, insertions, and deletions can be directly inspected within their local sequence context.

### PVG annotation visualization

Users can visualize annotation related to either one single genome or the complete PVG, and navigate through the different genes and their variations. Annotations can be provided as a sample-specific BED file (with position related to the current pivot), or as an ecumenic GAF file (such as the one provided through the PVG annotation by GrAnnoT [32]). Annotations are shown as blue thick lines on the corresponding nodes, solid when the current loaded annotation is identical to the pivot sequence (no variations), or split if variations occur (Figure 6).

**Fig 6.**
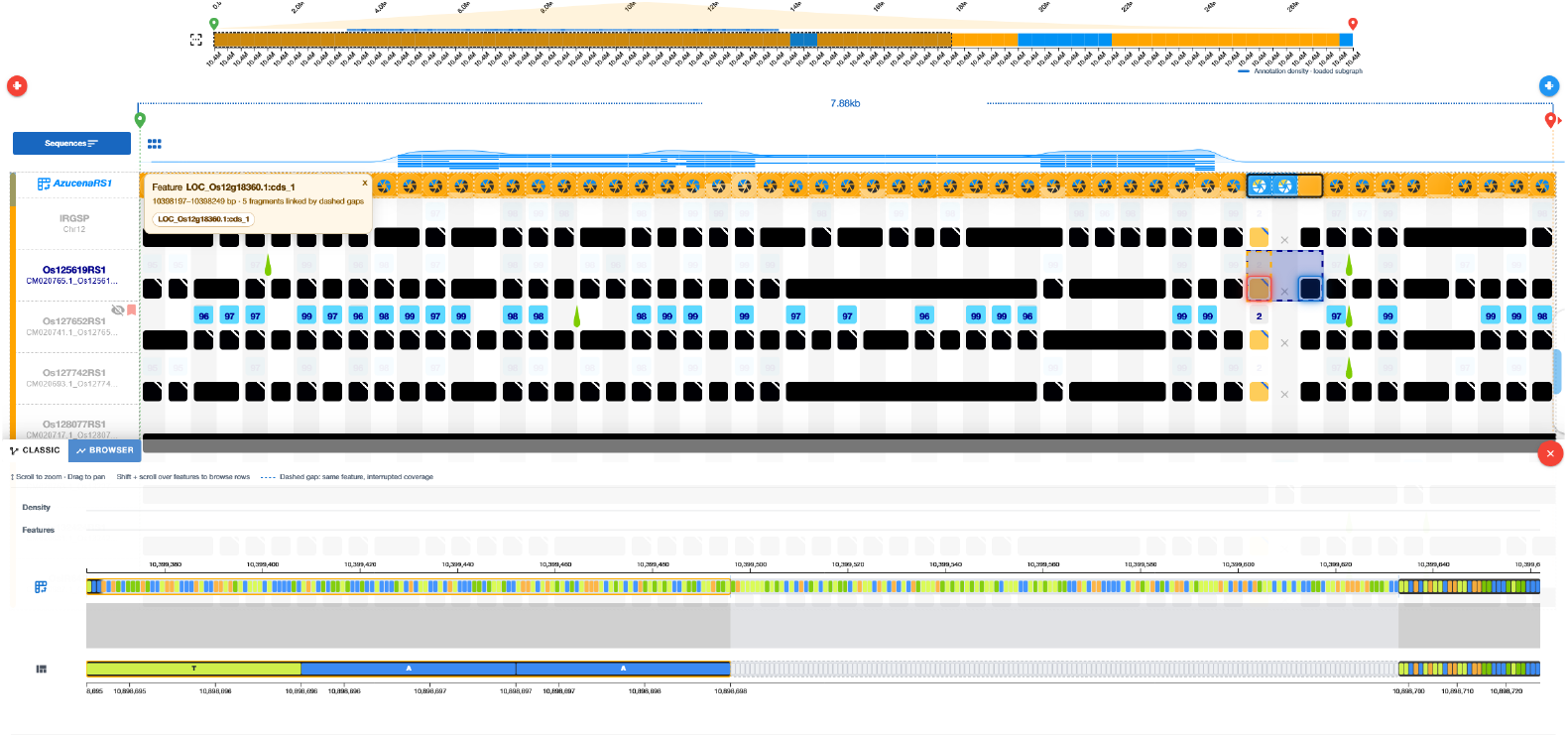
Visualization of the *Pi-ta* locus in Rice pangenome and associated structural variations. The annotation layer is displayed above the pivot genome, with annotated features represented by thick blue segments. The height of the thin blue profile indicates the proportion of genomes whose paths contain the corresponding annotated feature. If a feature is interrupted in the pivot genome, for example because of a deletion relative to the genome from which the annotation originates, the blue feature is displayed as discontinuous segments. At the *Pi-ta* locus (LOC_Os12g18360), comparison across genomes reveals structural variation affecting the region, including a deletion in a subset of genomes. The lower browser provides nucleotide-level inspection of the selected region and allows the underlying sequence differences to be examined in detail.

Once loaded, users can navigate either using the genomic positions of the annotations/genes they want to analyze, or directly through searching the genes/annotations names. They can thus directly identify the impact of variations on the different annotated features.

### Local graph view

Savanache also provides a more conventional, local graph-based representation of PVG data, using the pivot sequence as a baseline, conceptually similar to VRPG [26]. From the menu, users can select a region and generate a local graph centered on the corresponding pivot path (Figure 7). Alternative paths are displayed as branches around this baseline, allowing users to directly inspect the graph topology underlying the structural variation observed in the linear view. Nodes can be selected to link the graph representation back to their corresponding position in the structural variation view, while zooming and panning support exploration of more complex regions. To maintain interactive performance and readability, this representation is generated for user-defined local regions rather than for the complete pangenome graph .

**Fig 7.**
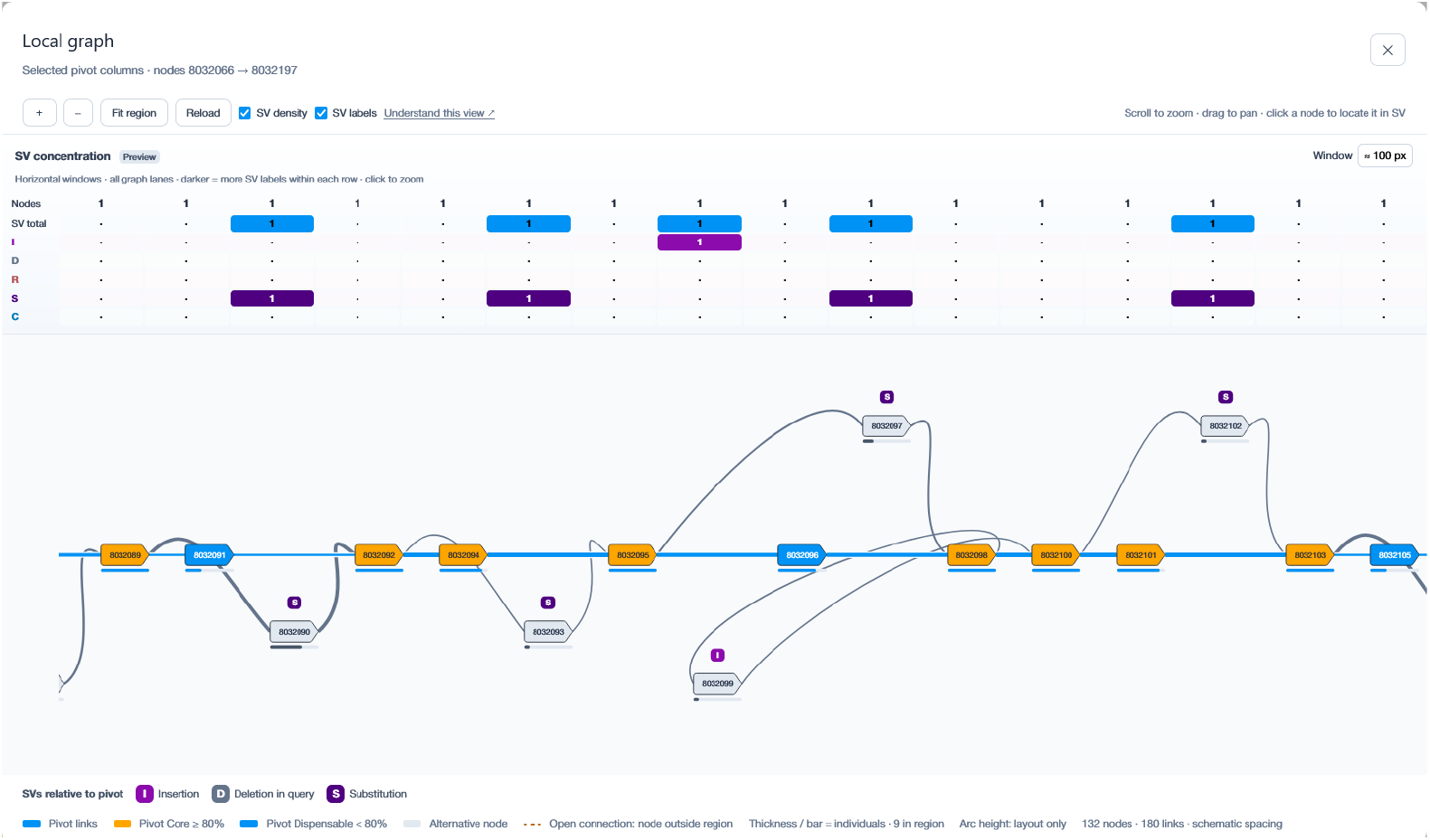
Local graph representation of a selected pangenome region. The local graph is generated from a user-defined region of the pivot genome. The pivot path is shown horizontally, while alternative graph paths are displayed as branches above or below it. Nodes are annotated according to their relationship with the pivot, including insertions (I), deletions (D), substitutions (S), reversals/inversions (R), and co-occurring events (C). Core (orange) and dispensable (blue) pivot nodes are distinguished according to the selected frequency threshold, whereas alternative nodes are shown separately from the pivot path. The upper summary panel reports the local concentration of structural-variation types across successive graph windows and can be used to navigate directly to regions of higher variation. Users can zoom, pan, and select individual nodes to locate the corresponding region in the structural variation view. Open connections indicate graph paths extending beyond the currently selected region.

### Multi-scale chromosome view

The upper view provides a genome-wide summary of the currently selected chromosome (Figure 3). Users can navigate by clicking on regions of interest or by entering genomic coordinates. Chromosomal blocks are shaded according to structural complexity; darker regions denote lower variation, while lighter shades reflect higher densities of rearrangements.

SaVanache provides a zoom system that adjusts the amount of detail shown in the graph. Users can set the minimum node size, where a value of 1 corresponds to nucleotide-level resolution. An optimal node size is automatically chosen when the page loads, but users can refine it as needed. When zoomed out, nodes smaller than the selected threshold are compressed in an artificial multinode (up to a minimal size at least identical to the threshold), while larger nodes remain uncompressed. For genomes other than the pivot, identical nodes are highlighted with a black box, while variant ones are colored. A color gradient indicates how similar the nodes are to one another, making it easier to spot conserved and variable regions within the compressed view.

The interface further supports interactive manipulation of genome tracks. Genomes can be reordered, grouped, or dragged according to user-defined criteria, such as phenotypic traits or population tags. This functionality enables users to visually explore potential associations between SVs and biological features carried by specific subsets of genomes.

Selected genomic regions can be exported through a dedicated interface. Users may download alignments or retrieve corresponding FASTA sequences via the *Get Fasta* option, enabling seamless transition from interactive exploration to downstream analyses.

## Discussion

Pangenome Variation Graphs are becoming the standard representation for eukaryotic pangenomes and are expected to play an increasingly central role in genome comparison analyses [6, 10, 33, 34]. However, this new standard comes with new data structures and concepts that are not yet fully supported by visualization tools, which were largely developed for linear genomes and synteny-based comparisons.

Although visualization of PVGs is a frequently requested capability, existing tools provide only partial solutions, either static or interactive [26, 35, 36]. However, they are all limited to a short region or very few individuals, or are non-interactive, static images (or both). Moreover, while graph-based visualizations exist, many scientists continue to prefer linear representations that align with established analytical practices and mental models.

These limitations motivated the development of SaVanache, a visualization framework designed to bridge the gap between PVG representations and intuitive exploration of genome diversity and of structural variations at chromosome level. Its design was shaped through extensive user feedback and iterative development, following best-practice recommendations for visualization tools [37], and involving both academic and private-sector collaborators and end-users.

Existing pangenome graph visualization tools provide complementary perspectives. Bandage [18] is particularly useful for inspecting local graph topology and sequence connectivity, whereas Panache [23] offers a linearized, block-based view of presence–absence variation across genomes. ODGI [20] provides a scalable command-line toolkit for graph manipulation and global visual summaries, and PanGraphViewer [22] enables coordinate- and annotation-guided inspection of local subgraphs. In contrast, SaVanache combines multi-scale exploration with a pivot-based, browser-oriented comparison of assemblies. Its structural variation views are designed to support the progressive examination of conserved and variable regions across multiple genomes, from chromosome-wide patterns to local pairwise comparisons. SaVanache should therefore be viewed as complementary to these tools rather than as a replacement for their respective graph-topology, graph-processing, or annotation-centred capabilities.

Users feedback consistently indicated a strong need for multi-scale exploration, ranging from global pangenome composition to chromosome-level and structural variation detail. SaVanache therefore adopts a hierarchical navigation strategy that enables progressive exploration, moving from high-level views of sampling and pangenome comparison to fine-scale inspection of genetic diversity and structural rearrangements.

The increasing number of genomes incorporated into PVGs raises important questions about sampling strategies, data set composition, genetic relationships, and representativeness. These considerations often motivate the construction of multiple pangenomes with diverse sampling schemes, leading to a growing need to compare pangenome contents and identify shared or overlapping individuals. This has been the rationale for the development of a dedicated page to explore and compare graph content together and enabling the description of individual genome assemblies used for constructing the graph.

While PVGs provide a powerful data structure for managing deconstructed genomic information across multiple references, their abstract nature can otherwise obscure biological interpretation. Users reported that they generally do not want to visualize the full graph topology directly: instead, they want to examine a genome of interest and compare it simultaneously with all others. To accommodate this expectation, SaVanache’s primary design principle is to represent all SVs within a PVG by maintaining a linear-like representation of individual genomes in its main view. Linear-based views are preserved to support intuitive navigation and interpretation, reducing visual complexity by emphasizing biologically meaningful patterns and relationships.

While the linear-like representation necessarily sacrifices some of the topological richness inherent to PVGs, enabling users to dynamically switch from a pivot genome to another preserves familiar workflows while leveraging the expressive power of PVGs, allowing users to focus on a chosen pivot genome without losing access to the full diversity encoded in the PVGs. In addition, the possibility to explore the same view in a more classical ‘graph’ vision using the pivot as main path allows a structural exploration for expert users.

Finally, the annotation layer, showing all the features located on a set of nodes, permits a very powerful analyses of biological insights of SVs and of their impact on genes. This annotation, either based on the current pivot (coordinates-based) or on the graph itself (coordinates-agnostic, see [32]), can help to explain major phenotypic variations between embedded individuals.

From a technical perspective, several key design choices support this goal. First, SaVanache employs an abstract segment representation, in which node lengths are standardized independently of their genomic size. This abstraction emphasizes structural relationships and organizational patterns rather than physical distance, improving the visibility of small but biologically significant elements, and facilitating comparison of SVs across different scales, particularly in regions with complex rearrangements.

Second, the interface is intentionally designed to limit the number of information layers displayed simultaneously. This choice minimizes cognitive load and prevents visual overload, which is especially important when navigating such complex graph-derived representations. While additional node-level information, such as gene annotations, may be incorporated in the future as small pop-up features, uncontrolled layering would compromise usability.

SaVanache also serves as a powerful diagnostic asset, as it faithfully reflects the underlying structure, qualities and defaults of the PVG and of its embedded assemblies. While the tool does not programmatically distinguish between real biological SVs and technical artifacts — such as those arising from misassemblies or alignment errors — it ensures these discrepancies are clearly visible within the interface. This visibility allows the multi-genome visualization to act as a natural validation layer: while SVs shared across multiple individuals reinforce their potential biological plausibility, the clear visibility of unique or irregular patterns empowers the user to intuitively identify and investigate potential technical artifacts. By exposing these details rather than obscuring them, SaVanache turns the complexity of pangenome data into a reliable opportunity for rigorous data validation.

Finally, real-time visualization of PVGs is computationally demanding. SaVanache addresses this challenge through a fast and efficient graph-indexing strategy optimized for interactive exploration of large PVGs (2h maximum - real time - for a 69 haplotypes graph). These data structures enable responsive operations such as pivot switching, zooming across scales, and dynamic updates of the visualization.

## Conclusion

SaVanache has been developed in close interaction with end-users to address a broad range of needs for the exploration of pangenomes. In addition to its visualization capabilities, it provides a large number of supporting resources, through videos or helping pop-ups. The public interface helps biologists to manipulate SaVanache before its local deployment. In terms of scalability, SaVanache already supports large PVGs, as the 69 *Arabidopsis thaliana* individuals available in the public instance. However it remains best suited to large screens for a good readability.

SaVanache is continuously under development and improvement, as we will add new features, while respecting our own rules [37], such as extraction of local haplotypes. In addition, we aim to facilitate the ingestion of new PVGs. Further optimization will also be required to support increasingly large datasets, such as the last version of the human pangenomes (a 10-haplotypes version of Chromosomes 1 to 7 is available as an example on the public interface). Overall, SaVanache provides an interactive framework to navigate the complexity of graph-based pangenomes and exploring structural variation across multiple genomes.

## Supporting information

SuppTable 1

SuppFigure 1

## Supporting information

**S1 Table. Comparative table of pangenome visualization tools’ representations and data** Comparison of visualization paradigms, visual abstractions and scales, and input data for existing pangenome visualization tools.

**S1 Figure Download of sequence alignments from the Structural Variation Browser**. Users can select a genomic region directly in the Structural Variation Browser and download the corresponding sequence alignment using the “download region >Get Align” button on the right side of the interface or advanced menu. The exported file contains multiple pairwise CLUSTAL W like alignments between the selected pivot genome and each of the other genomes for the selected region.

## Design and implementation

All codes used in the current version (1.0.1) are available at the IRD DataSuds repository for Savanache for the stable version and at the Savanache main IRD Forge repository for the development version.

### Data collection

A set of publicly available pangenome datasets was identified across different species to populate and test SaVanache [32]. For datasets where a pangenome variation graph (PVG) was not already available, one was constructed using MiniGraph-Cactus, following the standard ‘–gfa full’ output format required by SaVanache (see GitLab documentation for format specifications), in the same way as in Marthe et al [32]. All resulting GFA files, whether pre-existing or newly generated, were subsequently indexed using the indexing pipeline described in the SaVanache documentation, which builds the structural indexes, SV cache, duplication index, and Mash distance matrices (see Section Diversity) required for interactive visualization.

### Interactive visualization (Frontend)

SaVanache is written in JavaScript. It is part of the tools from the GraSuite (https://forge.ird.fr/diade/grasuite), licensed through GNU GPLv3 and available on the IRD Forge (https://forge.ird.fr/diade/savanache). A Docker compose file allows an easy deployment, and a public instance is available at https://savanache.ird.fr.

Its front-end is built using the JavaScript framework **Vue.js 3** and related dependencies ( **Vuex** for storing variable states across components; **Vue Router** for client-side navigation between pages; **Vuetify 3** for design elements). Data visualization is handled with the libraries **amCharts 5** and **D3.js**. The interface is structured around four main views, corresponding to the different pages of the application.

Specific dynamic interactions are implemented using **jQuery**, particularly for targeted DOM manipulations.

Communication with the backend is performed via **Axios** for HTTP requests, and through the standard **WebSocket** API for real-time synchronous exchanges.

### Technical implementation (Backend)

The backend is written in Python, using **Django v5**, and consists of two applications: one in WSGI (synchronous) mode and the second in ASGI (asynchronous) mode. The latter utilizes the SaVanache API, which manages indexing to optimize data access speed and SV calculations, while providing metadata related to visualizations. The API consists of three modules: a dedicated PVG parser tool for managing indexes and data formats, a module for managing the SQLite database, and a module handling graph navigation that communicates with both the indexes and the database.

In practice, the parsing tool functions as a PVG indexing tool designed to provide a fast user experience when accessing graph data. It currently supports GFA spec version v1.1 and focuses only on the Segments (S), Links (L) and Walks (W) lines of the GFA file.

Segments (S) lines represent the sub-sequences belonging to the individuals of the graph, and links (L) lines represent adjacency between two segments. Finally, the ordered succession of S and their orientation (forward or reverse-complement) represents walks (W), the succession of segments orientation for a specific individual in the graph.

The parser uses the specific GFA 1.1 output structure from Minigraph-Cactus (version v2.8+, [33]). In this format, the Segments are ranked by ascending order of IDs, which are natural numbers. The Walks are grouped by chromosome, and the Links are ranked by minimal segments identifiers that they link. For each pair of *S* ×*S* ∈ *L*, the ranking is based on *min*(*S*_*a*_, *S*_*b*_). The set of *L* is ordered in ascending order of the value min(*a, b*) for each pair (*a, b*) ∈ *L*, where *a* and *b* denote the identifiers of the connected S’s.

Users can convert PGGB GFA files [10] into the required format using GraMER (https://forge.ird.fr/diade/graphgwas/gramer) before importing them into SaVanache.

### Parser tool Structures

To optimize data extraction from GFA files, SaVanache generates a set of index and metadata files (Figure 8). The system creates index files for both Segments and Walks, enabling *O*(1) time complexity for direct access to these elements within the GFA file. Additional metadata, binary files store the restructured GFA information and pre-computed metrics, providing access times ranging from *O*(1) to *O*(*logn*), where *n* represents the volume of stored data. The system also maintains a relational database containing the step (see below) of each segment in each walk for each chromosome, to facilitate fast query processing.

**Fig 8.**
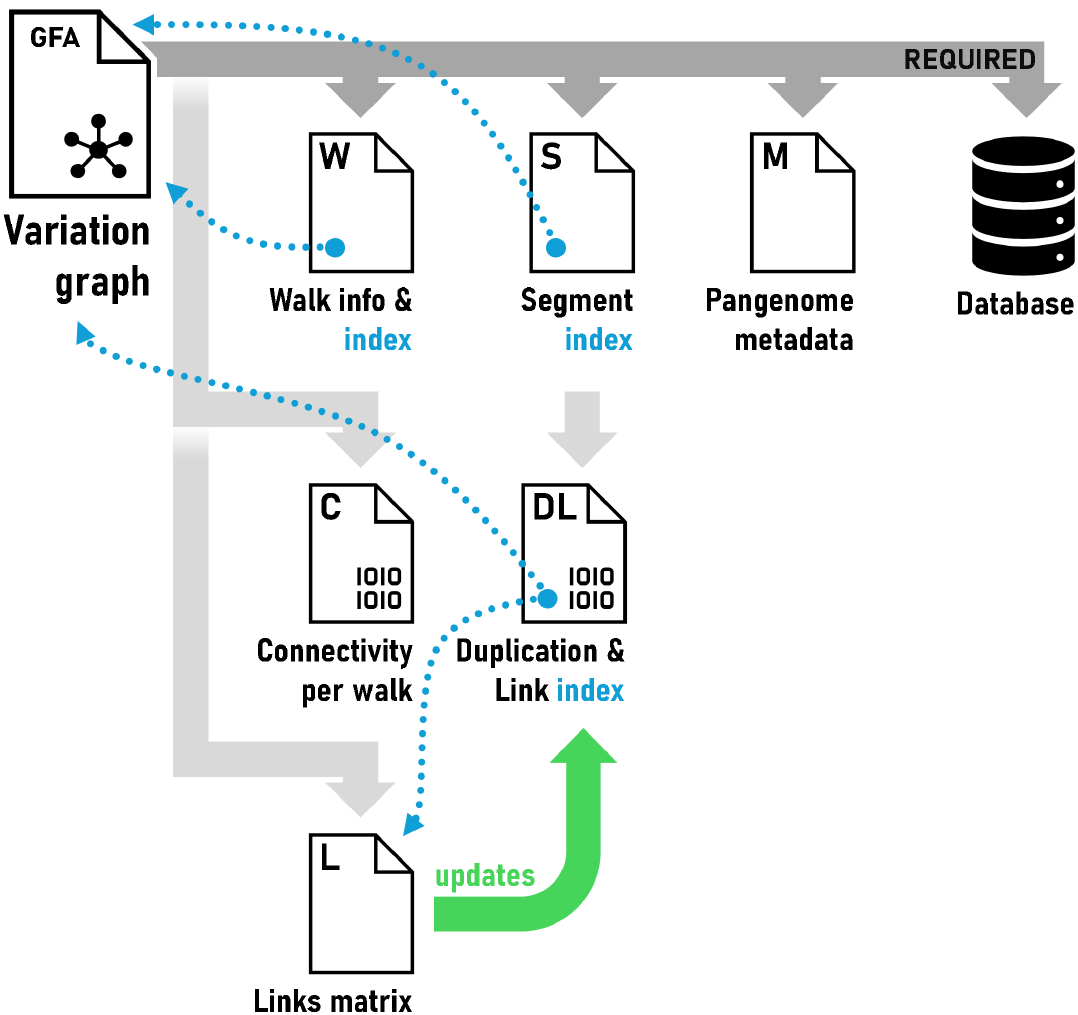
Files derived from the original GFA during SaVanache preprocessing. The minimum set of required files to run a successful SaVanache session is created first: two index files—one for the Walks (***W*** ) and the second for Segments (***S***)—complete with metadata for the whole pangenome (***M*** ), and a relational database with positional information of segments. For query optimization purposes, two additional binary files are created. The first extracts the actual connectivity of Segments per Walk (***C***) from the GFA. The second is a compression of ***S*** updated down the line with the index of extracted Links (***L***), effectively tracking inter/intra-chromosome duplications (***DL***).

### Positions and Steps

SaVanache uses Walk information from the GFA file to identify sequence conservation and SVs between individuals embedded in the PVG. We introduce two key concepts related to Segments within a Walk, the *Position* and the *Step*:

- *Position* refers to the genomic coordinates (chromosome and base-pair position) of a given Segment (biological context),
- *Step* represents the sequential index of a Segment within the ordered list of Segments defining the Walk (informatic context).

### Walks binary format

The Walks from the GFA file are represented in a binary format, allowing direct access to their *Steps* and *Positions*. The data is stored in the `<graph.gfa>.idx_bin` file, while an associated index, `<graph.gfa>.idx_bin_idx.json`, retains the metadata necessary to locate the Walks and their binary lines.

Let *C* be the maximum number of Segments stored in a binary line, *R* the number of lines allocated to a Walk, *B*_*n*_ the number of bytes required to encode a Segment identifier, and *B*_*p*_ the number of bytes used to represent a nucleotide coordinate. A Walk *w* containing *n*_*w*_ Steps is represented by a block consisting of a header of *B*_*n*_ bytes, containing *n*_*w*_, followed by *R* fixed-size lines.

A line *r* encodes the nucleotide interval [*b*_*r*_, *e*_*r*_) it covers, followed by *C* segment cells and an end-of-line marker. Its size is thus:

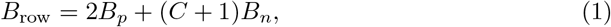

where *b*_*r*_ and *e*_*r*_, each encoded using *B*_*p*_ bytes, correspond respectively to the start and end nucleotide coordinates of the line. Each segment cell, as well as the end-of-line marker, occupies *B*_*n*_ bytes.

The size of a block representing a Walk is therefore:

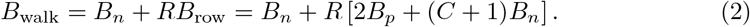

Since the parameters *C, R, B*_*n*_, and *B*_*p*_ are fixed during the construction of the index, the blocks associated with the Walks have a constant size. The position of a Walk in the binary file can thus be calculated directly from its identifier, without traversing the preceding Walks.

The layout of a block is illustrated in Fig. 9. Denoting *s*_*r,c*_ as the *c*-th segment cell of row *r*, the content of a Walk can be summarized by:

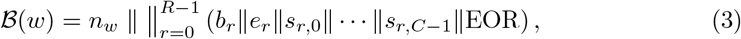

where ∥ denotes the concatenation of binary representations and EOR the end-of-line marker.

**Fig 9.**
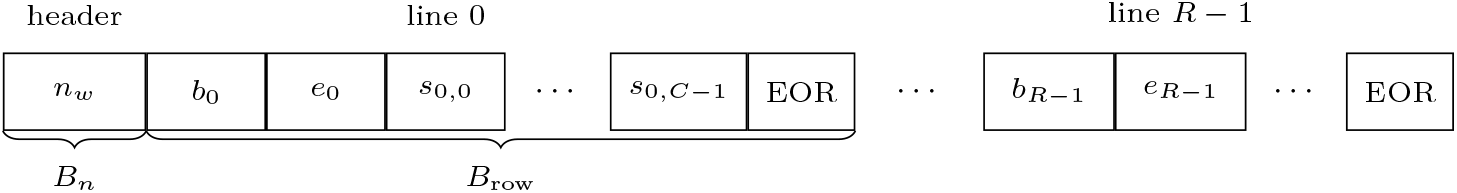
Organization of the binary format of a Walk. Each Walk is represented by a header containing its number of Steps, followed by *R* fixed-size rows. Each row contains its nucleotide boundaries, *C* Segment cells, and an end-of-line marker.

A cell *s*_*r,c*_ simultaneously encodes the Segment identifier and its orientation in the Walk. For a cell of *B*_*n*_ bytes, the most significant bit represents the orientation and the remaining 8*B*_*n*_−1 bits represent the Segment identifier. If *x* denotes the unsigned integer value of a cell, the corresponding identifier is:

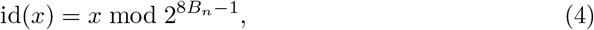

and its orientation is given by:

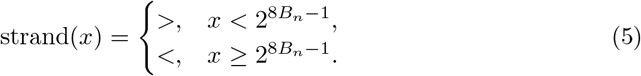

The index file also associates with each row *r* of the Walk *w* the nucleotide interval [*b*_*r*_, *e*_*r*_). These intervals make it possible to locate a region of the Walk from nucleotide coordinates without storing the individual position of each *Step*. The lengths of the Segments are not duplicated in this representation and are obtained from the Segment index during navigation.

#### Spatial complexity

The width of the binary fields is not fixed independently of the PVG. It is determined before the construction of the index based on the largest values to be represented. Let *I*_max_ be the maximum value of a Segment identifier and *P*_max_ be the largest nucleotide coordinate. The size *B*_*n*_ of a Segment cell is chosen as the smallest number of bytes allowing the encoding of *I*_max_ while reserving the most significant bit for the orientation. It thus satisfies:

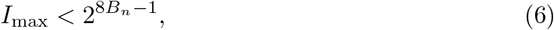

and can be expressed by:

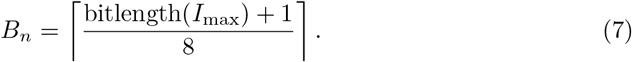

Similarly, the size *B*_*p*_ used for nucleotide coordinates corresponds to the smallest number of bytes allowing the encoding of *P*_max_,

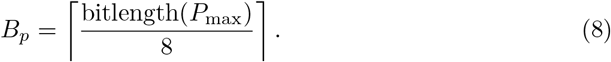

For *N*_*w*_ walks, the total size of the binary representation is therefore

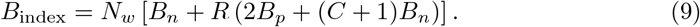

The space complexity can thus be expressed by:

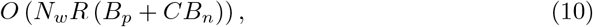

or, by making explicit the dependence of the representation width on the maximum values of the pangenome:

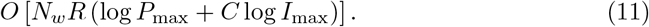

This representation favors direct data access at the expense of compactness. In particular, the widths *B*_*n*_ and *B*_*p*_ are determined by the largest values present in the pangenome and then used uniformly throughout the index. Increasing the size of the pangenome can therefore increase not only the number of cells to be stored, but also the individual size of each of these cells.

### Auxiliary segment indexes

In addition to the binary representation of Walks, we maintain auxiliary structures associated with the graph segments. In particular, they allow direct access to Segment properties, the identification of sequences shared by multiple identifiers, and, optionally, the representation of adjacency between Segments.

#### Segment index and sequence duplication

Each numerical Segment is associated with a fixed-size record in the file `<graph.gfa>.idx.p.s.bin`. A record contains the information necessary to access the Segment as well as auxiliary fields for duplication and links:

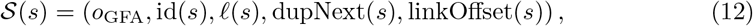

where *o*_GFA_ denotes the position of the Segment in the GFA file, *l*(*s*) its nucleotide length, dupNext(*s*) a pointer to a Segment carrying the same sequence, and linkOffset(*s*) a pointer to the auxiliary Link index.

These five fields use a fixed width of *B*_*s*_ bytes, determined from the largest value to be represented. The record of Segment *s* can thus be directly located according to:

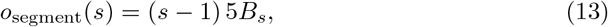

In particular, it distinguishes Segments that have different identifiers but carry exactly the same sequence. For two Segments *s*_*i*_ and *s*_*j*_:

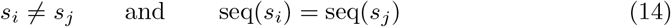

define an exact sequence duplication. Segments belonging to the same sequence class are linked by the dupNext field in the form of a circular chain. A Segment can thus be identified as belonging to a duplicated class when:

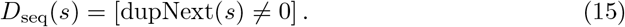

This information characterizes an exact sequence equality between multiple identifiers and must be distinguished from the repetition of the same identifier within a Walk, which is detected from the position database described in Section *Locating a corresponding region in a Walk*.

#### Auxiliary Link index

We also features an optional topological index built from the L lines of the GFA. For a Link between two Segments *u* and *v*, the current implementation defines a normalized representation:

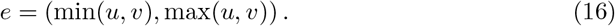

Links are grouped by chromosome in a file

`<graph.gfa>.link.diversity.char.<chrom_id>`.

For a Link *e* and a Walk *w*, the index stores a value:

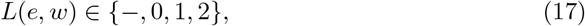

where 0 indicates that the adjacency is absent from the Walk, while 1 and 2 indicate that it is observed in the two possible orders of the normalized identifiers. The orientation carried by the +/-fields of the GFA is not directly represented by this value.

The linkOffset(*s*) field of the Segment index points to the first Link entry associated with Segment *s*. If *d*(*s*) denotes the number of normalized adjacencies starting with *s*, their retrieval can be performed in:

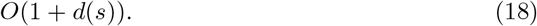

The index thus allows determining which Walks traverse a given adjacency between two Segments.

In the current implementation, this topological index remains experimental and does not directly intervene in the structural variation detection pipeline described in the following sections. It nevertheless constitutes an additional source of information that could be used to disambiguate repeated occurrences based on their topological context.

### Access and navigation in Walks

The regular structure of the binary format allows accessing Walks either from the position of a *Step*, or from a nucleotide coordinate. Let *n* be the number of *Steps* contained in a Walk and *C* the maximum number of Segments that can be stored in a binary line. The position of a *Step* in the file can be determined directly, while searching by nucleotide coordinate relies on the intervals associated with the binary lines.

#### Direct access by *Step*

Let *s*, with 0 ≤ *s < n*, be the index of a *Step* in a Walk. The row *r* and column *c* containing this *Step* are obtained directly by:

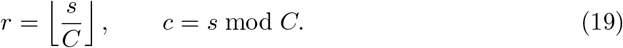

Denoting *o*_chr_ as the starting position of the chromosome in the binary file and *w* as the identifier of the Walk within it, the address of the cell corresponding to *Step s* is:

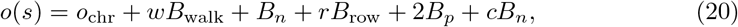

where *B*_walk_ and *B*_row_ correspond respectively to the sizes of a Walk block and a binary row defined in Section *Direct access by Step*. The term *B*_*n*_ allows bypassing the Walk header and 2*B*_*p*_ the nucleotide coordinates stored at the beginning of the row.

The Segment and its orientation are thus obtained by directly positioning the read cursor at address *o*(*s*) and then reading *B*_*n*_ bytes. No traversal of the *Steps* preceding *s* is necessary. The time complexity of accessing a *Step* is therefore:

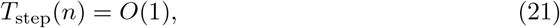

with constant auxiliary memory *O*(1). This property is independent of the number of *Steps* present in the Walk.

#### Extraction of a *Step* interval

Direct access can be applied successively to an interval of *k Steps*. For an interval [*s*_start_, *s*_end_], with:

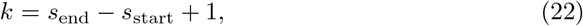

each *step* is obtained from its address calculated according to Equation 20. The extraction consequently requires:

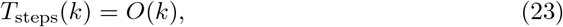

and the memory required to represent the result is *O*(*k*). The complexity thus depends solely on the number of Segments actually returned and not on the total length of the Walk.

#### Conversion of a *Step* to a nucleotide coordinate

The calculation of Equation 19 also allows directly locating the row containing a *Step s*. However, the binary format does not store the individual nucleotide coordinate of each Segment. Only the coordinate *b*_*r*_ corresponding to the beginning of row *r* is recorded.

Let *l*(*s*_*r,j*_) be the nucleotide length of the Segment stored in the *j*-th cell of row *r*. The nucleotide coordinate of the beginning of the Segment associated with *Step s* is then:

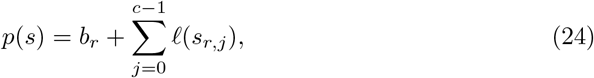

where *c* = *s* mod *C*. The lengths *l* are obtained from the Segment index and are not duplicated in the binary format of the Walks.

In the worst case, the searched Segment occupies the last cell of the row and up to *C −* 1 lengths must be summed. The time complexity of this conversion is therefore:

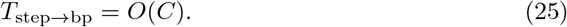

In the current implementation, *C* is bounded to 10, 000 Segments. The cost of this operation is therefore independent of *n* when this bound is considered fixed.

#### Access by nucleotide coordinate

The inverse operation consists of determining the Segment containing a nucleotide coordinate *p*. The metadata file associates with each row *r* an interval:

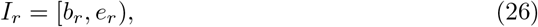

whose end bounds *e*_*r*_ are ordered according to their position in the Walk. If *R*_*n*_ denotes the number of rows actually used to represent the *n Steps* of the Walk, then:

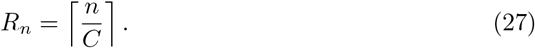

The search consists of determining the first interval whose upper bound is strictly greater than the searched coordinate:

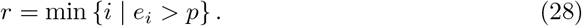

The obtained interval indeed contains the coordinate when:

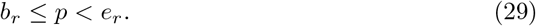

Since the bounds *e*_*r*_ are ordered, *r* can be determined by binary search among the *R*_*n*_ intervals. This operation requires:

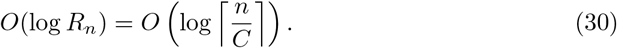

Once the row is identified, the lengths of the Segments it contains are accumulated from *b*_*r*_ until reaching the coordinate *p*. At most *C* Segments must be examined. The worst-case complexity of locating a Segment from a nucleotide coordinate is thus:

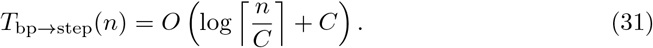

Since *C* is bounded independently of the Walk length, it can be considered constant in the asymptotic analysis. We then obtain:

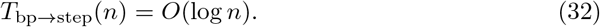

This organization thus reduces the search for a nucleotide coordinate to a search among approximately *n/C* binary rows, followed by a local traversal bounded to *C* Segments, rather than a traversal of the *n Steps* of the Walk.

#### Extraction of a nucleotide region

For a nucleotide region [*p*_start_, *p*_end_), the first row is located according to the preceding procedure. The Segments are then read sequentially until reaching *p*_end_. If *k* Segments overlap the requested region, the extraction cost becomes:

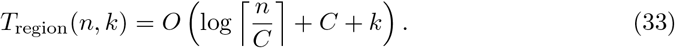

When *C* is considered constant, this expression reduces to:

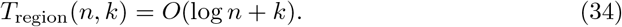

### Locating a corresponding region in a Walk

After extracting a pivot region, we search in each query Walk for a region likely to correspond to it. This localization relies first on searching for occurrences of shared Segments, and then on chaining candidate anchors to identify a structurally coherent region.

#### Search for candidate occurrences

For each chromosome, a SQLite table associates Segment identifiers with their occurrences in the different Walks. An occurrence is represented by:

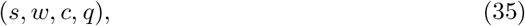

where *s* is the Segment identifier, *w* the Walk, *c* the chromosome, and *q* its position expressed in *Steps*. For a set *S* of Segments from one end of the pivot region and a query Walk *w*_*q*_, the search can be written as:

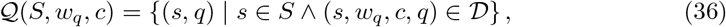

where D denotes the set of occurrences recorded in the database.

The pivot Segments are queried in successive windows from each of its ends until a sufficient number of distinct Segments present in the query Walk is obtained. If *M* denotes the number of occurrences indexed in the considered table, *b* the number of queried Segments, and *H* the number of returned occurrences, the cost of this step can be approximated by:

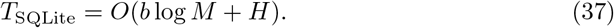

#### Individual weight of anchors

Each shared occurrence defines a candidate anchor:

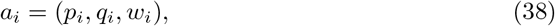

where *p*_*i*_ and *q*_*i*_ correspond respectively to the positions of the Segment in the pivot window and in the query Walk. The weight *w*_*i*_ estimates the confidence granted to the anchor and combines the Segment length, its multiplicity in the pivot and the query, its potential status as a duplicated Segment, and its proximity to the searched edge:

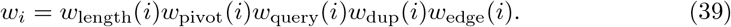

The component related to length is defined by:

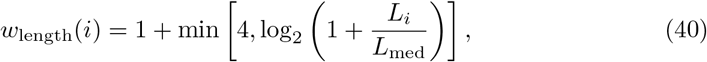

where *L*_*i*_ is the length of the Segment and *L*_med_ the median length of the Segments in the window. Repetitions are penalized according to:

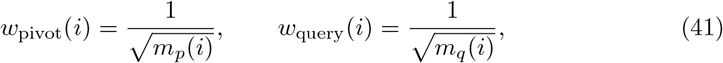

where *m*_*p*_(*i*) and *m*_*q*_(*i*) represent the number of occurrences of the Segment in the pivot and in the query. Segments explicitly marked as duplicated also receive a factor *w*_dup_ = 0.25.

For a window of *b* Segments, the edge proximity factor is defined, for the search of the beginning and the end respectively, by:

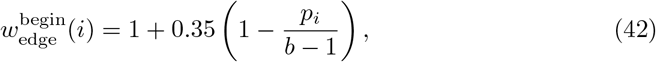

and :

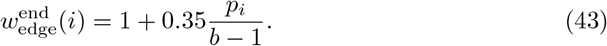

#### Anchor chaining

The anchors are then chained by dynamic programming. They are considered in increasing order of their position in the pivot, so that a transition from *a*_*j*_ to *a*_*i*_ requires:

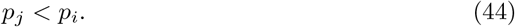

Two orientations *o* ∈ +1, −1 are evaluated separately, and a transition is admissible when

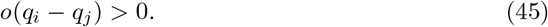

The orientation *o* = +1 corresponds to a collinear organization of the anchors, while *o* = −1 allows searching for an inverted organization.

The structural coherence of two anchors is evaluated from the number of *Steps* separating them:

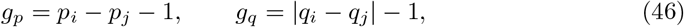

and their normalized distortion:

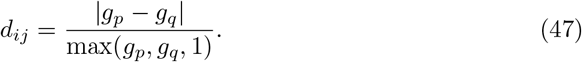

A valid transition receives a proximity bonus:

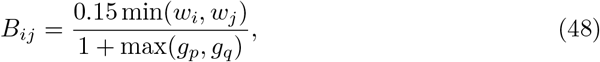

and a distortion penalty:

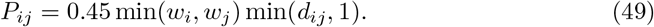

For an orientation *o*, the best score of a chain ending at anchor *a*_*i*_ is then:

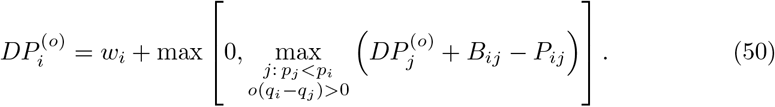

This formulation favors chains composed of reliable and structurally close anchors, while tolerating spacing differences likely to correspond to variations between Walks.

#### Chain selection

A reconstructed chain finally receives a bonus based on its distance *D*_edge_ to the searched edge:

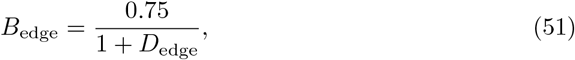

as well as a penalty when it contains fewer than three anchors:

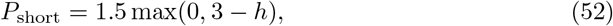

where *h* is the number of anchors in the chain. Its final score is:

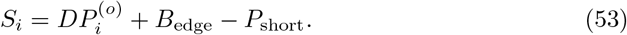

The best chain between the forward and reverse orientations is kept. The coefficients involved in the score are empirical hyper-parameters chosen to favor structural coherence and support by multiple anchors without excluding regions with significant variations.

#### Complexity

Let *A* be the number of candidate anchors. Sorting them requires *O*(*A* log *A*), while dynamic programming compares, in the worst case, each anchor to the anchors preceding it. The complexity of `find_best_anchor()`is thus:

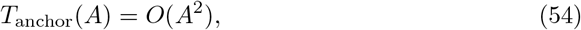

with *O*(*A*) auxiliary memory. In the current implementation, the number of candidates is limited by the size of the queried windows and by the maximum number of occurrences kept for each segment.

### Construction of shared anchors between sub-Walks

Once the query region is localized and extracted, we establish correspondences between occurrences of shared Segments in the pivot and query sub-Walks. Unlike the previous step, which aims to globally localize the query region, this step seeks to precisely associate homologous occurrences of the Segments present in both regions.

For a segment *x*, let:

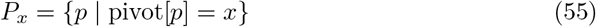

and:

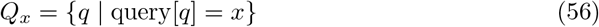

be the sets of its local positions in the pivot and in the query. Only Segments such that *P*_*x*_*≠* ∅ and *Q*_*x*_*≠* ∅ are kept.

#### Unambiguous anchors

When a Segment appears only once in each of the two sub-Walks:

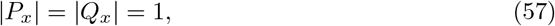

its correspondence is directly defined by:

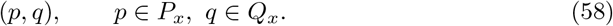

These correspondences constitute a set of unambiguous anchors U. No global monotonicity constraint is imposed on their positions in the query. Increasing or decreasing sequences of anchors can thus be preserved and used later to characterize collinear or inverted regions.

#### Resolution of repeated segments

When a Segment appears multiple times in at least one of the two sub-Walks, the set:

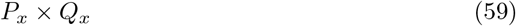

defines several possible correspondences. we then use neighboring unambiguous anchors as a local context to select the most coherent pairs.

For a pivot occurrence *p*, let:

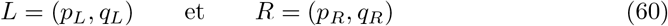

be the nearest unambiguous anchors located to the left and right of *p*, respectively. When both flanks are available, the local orientation is defined by:

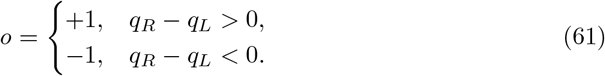

A candidate correspondence (*p, q*) must be compatible with this local framework. For *o≠* +1, the query position must satisfy:

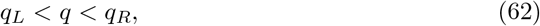

while for *o* = −1:

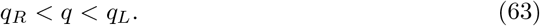

This formulation allows collinear and inverted contexts to be treated identically.

#### Contextual score

The coherence of a candidate pair (*p, q*) is evaluated from neighboring unique anchors. For a context anchor *a≠* (*p*_*a*_, *q*_*a*_) *∈ U*, we define:

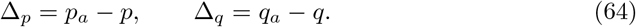

The anchor is compatible with the local orientation when:

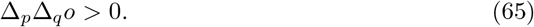

The associated structural distance and distortion are, respectively:

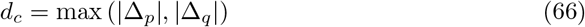

and:

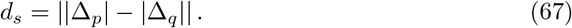

A context anchor contributes to the score when *d*_*c*_≤ 8 and *d*_*s*_ ≤2. Its contribution is then:

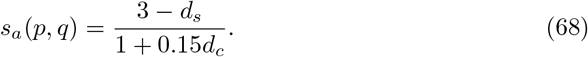

The contextual score of a candidate correspondence is defined by:

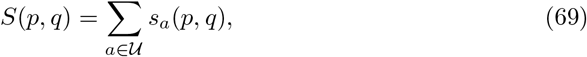

where only anchors satisfying the preceding conditions contribute to the sum. This score favors correspondences supported by nearby anchors and exhibiting similar spacing in both sub-Walks.

Candidates are then ordered by their contextual score, their number of supporting anchors, and their local distortion. They are greedily selected under the constraints:

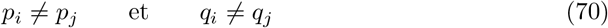

for two distinct selected correspondences. Each occurrence of a repeated Segment can thus be associated with only a single occurrence in the other sub-Walk.

#### Fully repetitive regions

When no unambiguous anchor is available, no local context can differentiate the various occurrences of a repeated Segment. We then use a conservative matching based on their order:

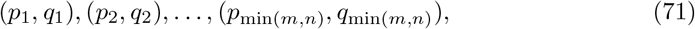

where *m≠* |*P*_*x*_| and *n≠* |*Q*_*x*_ |. This mechanism acts as a fallback solution for fully repetitive regions.

The selected correspondences are finally represented in the form:

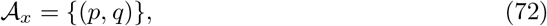

for each shared Segment *x*. They are serialized by save_node_shared and used later during the comparative traversal of the two sub-Walks.

#### Ambiguity of correspondences

The presence of multiple possible occurrences of a Segment is not the only source of ambiguity when constructing shared anchors. We distinguish several complementary signals.

The first corresponds to Segments with distinct identifiers but a strictly identical sequence, detected by the segment index:

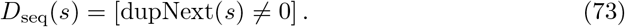

The second corresponds to the local repetition of the same identifier in the two compared sub-Walks:

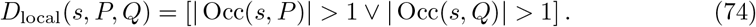

The position database can also indicate that the same identifier appears multiple times in a full Walk:

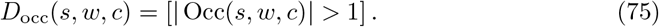

These signals describe different phenomena: *D*_seq_ indicates that the same sequence can be represented by multiple identifiers, *D*_occ_ that an identifier is repeated within a Walk, and *D*_local_ that an ambiguity is indeed present in the two regions currently being compared.

In the representation used by save_node_shared, a Segment is marked as ambiguous when at least one of these signals implies multiple possible correspondences. This information is preserved in the DUP field and propagated to structural events via an additional annotation. It does not alter their primary class, but indicates that their interpretation relies on a potentially ambiguous correspondence.

#### Complexity

Let *N*_*P*_ and *N*_*Q*_ be the number of *Steps* in the pivot and query sub-Walks, *U≠* |*U*| the number of unambiguous anchors, and:

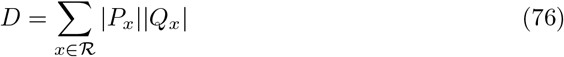

the total number of candidate correspondences for repeated Segments *R*. Building the occurrence sets requires:

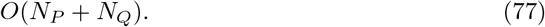

In the current implementation, evaluating a candidate can traverse the *U* unambiguous anchors to calculate its contextual support, representing a cost of:

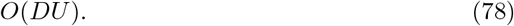

Sorting the *D* candidates furthermore requires:

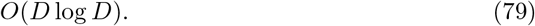

The total time complexity of constructing the shared anchors is therefore, in the worst case:

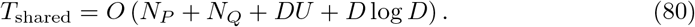

In weakly repetitive regions, *D* remains limited and the construction is close to a linear traversal of the two sub-Walks. Conversely, the cost can increase in highly repetitive regions due to the number of potential pairs |*P*_*x*_||*Q*_*x*_|.

### Structural Variation Detection and Classification

Once the correspondences between shared occurrences are established, the system performs pairwise comparisons between the pivot sub-Walk:

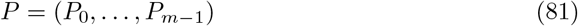

and query sub-Walk:

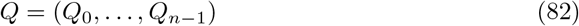

to identify structural differences. This process relies on the set of shared anchors:

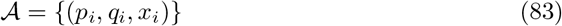

where *p*_*i*_ and *q*_*i*_ denote the local positions of the same Segment occurrence *x*_*i*_ in *P* and *Q* respectively.

#### Identification of inverted block

The anchors are first sorted according to their position in the pivot. The algorithm distinguishes inverted regions from collinear regions by evaluating the relative order of anchors. A sequence of anchors forms an Inversion (Inv) block when:

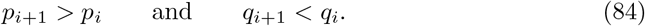

For such blocks, the internal intervals *G*_*P*_ and *G*_*Q*_ are compared by considering *G*_*Q*_ in reverse order to match the block’s orientation. Multiple consecutive inverted Segments are classified as chained inversions. In these cases, if both intervals are non-empty, the variation is classified as a SWAP regardless of length differences, and no residual insertion or deletion is inferred.

#### Event Classification and Labeling

Outside of inverted blocks, shared anchors delimit compared intervals *V*_*P*_ and *V*_*Q*_ in both sub-Walks. The variation between two successive anchors (*p*_*i*_, *q*_*i*_) and (*p*_*j*_, *q*_*j*_) is classified according to the occupancy of these intervals, as illustrated in Figure 10 and following these rules:

**Fig 10.**
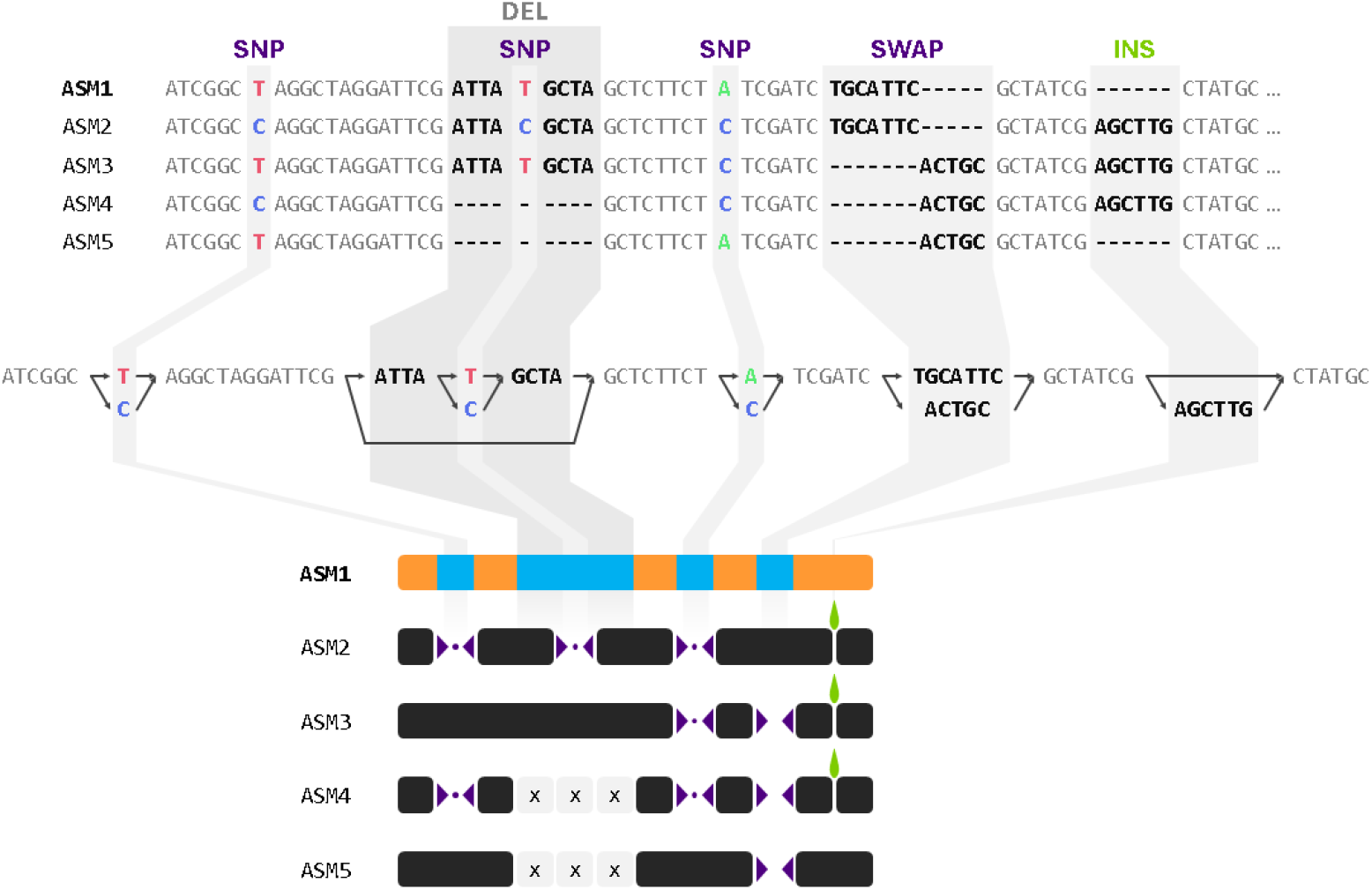
Equivalent representations of structural variations across multiple genomes in different representations. From top to bottom: multiple sequence alignement (MSA), Pangenome Variation Graph (PVG), SaVanache’s variation glyphs. The top panel shows a multiple sequence alignment of five individuals (ASM1 to ASM5) containing several types of variation: single nucleotide polymorphisms (SNPs), insertions (INS), deletions (DEL), and swaps (SWAP), where both pivot and query contain different sequences. The middle panel illustrates the same sequences visualized as a Pangenome Variation Graph (PVG), where conserved sequences are represented as nodes (segments) connected by edges (links). The bottom panel illustrates how such a PVG would look like when converted into SaVanache’s variation glyph representation, using ASM1 as the pivot assembly. In this representation, query assemblies (ASM2-5) are aligned to the pivot, with each structural variant classified and displayed as a specific glyph (for more detail on the glyphs please refer to section: *Glyph-based representation of SVs* in the Results section). On the ASM1 track, orange segments represent regions shared across all genomes (core genome), whereas blue segments indicate regions present in only a subset of genomes (dispensable genome).

- Insertion (INS): When |*V*_*P*_ |*≠* 0 and |*V*_*Q*_|*>*0. The segments in *V*_*Q*_ are present only in the query relative to the pivot. Biologically: the pivot contains two consecutive syntenic Segments while the query contains one or more intermediate Segments between them.
- Deletion (DEL): When |*V*_*P*_ | *>*0 and |*V*_*Q*_ |*≠* 0. The segments in *V*_*P*_ are present only in the pivot. Biologically: the pivot contains one or more intermediate segments between syntenic segments while the query contains no intermediate segments.
- Swap (SWAP): When both |*V*_*P*_ | *>*0 and |*V*_*Q*_ | *>*0, indicating a structural replacement where both pivot and query contain different intermediate segments. Biologically: both pivot and query contain different intermediate segments between syntenic segments.
- Single Nucleotide Polymorphism (SNP): A specific case of SWAP where both *V*_*P*_ and *V*_*Q*_ contain only one intermediate segment of length 1. Biologically: both pivot and query contain only one intermediate segment of length 1.
- Match (M) When both intervals are empty (|*V*_*P*_ |*≠* 0, |*V*_*Q*_ |*≠* 0), the anchors are adjacent, and the segments are classified as a match if they share the same orientation. Biologically: both pivot and query have the same consecutive Segments. A single-point strand difference at an anchor is preserved as a local inversion event, distinct from the multi-anchor inverted blocks described above.
- Inversion (INV): When segments in the query show different orientation compared to their corresponding segments in the pivot. Multiple consecutive inverted segments are classified as chained inversions.

#### Ambiguity and Duplication Signals

SV detection is performed locally, making it dependent on the identified syntenic segments. Consequently, the reliability of detection increases with the width of the selected region and the number of syntenic Segments it contains.

To handle complex graph topologies, events are annotated with signals that indicate potential interpretative ambiguity without altering their primary class:

- **Sequence Duplication (***D*_*seq*_): Detected via the segment index when different identifiers carry identical sequences.
- **Local/Global Repetition (***D*_*local*_, *D*_*occ*_): Identified when a segment appears multiple times within the compared regions or the full walk.
- **Co-occurrence (COOC)**: An additional signal for potential duplication or inter-chromosomal translocation, assessed by checking if a segment appears multiple times within the database (intra-chromosomal) or the segment index (inter-chromosomal).

#### Traversal Progression and Complexity

After processing an interval, the used anchors are consumed, and the current positions in both sub-Walks advance. The suffixes remaining after the last pair of anchors are treated using the same INS/DEL/SWAP rules.

Let *A* be the total number of shared anchor occurrences and *V* the number of elementary events produced. The overall time complexity for this comparison is:

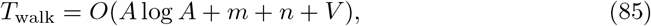

Since the number of produced events is normally bounded by the number of positions traversed, the traversal following anchor sorting is essentially linear in size.

### Recovery of structural variations

#### Detection of shared syntenic segments

When users define a region of interest by specifying genomic coordinates, the system identifies the two segments in the pivot assembly that bound this region based on their *Positions, i*.*e*. physical start and end coordinates on chromosomes. All Segments between these boundaries are extracted to form the pivot sub-Walk (Wp), with each Segment assigned to its *Step* index (representing its rank within the ordered segment list).

To extract corresponding sub-Walks from query assemblies, *i*.*e*., assemblies compared with the pivot assembly, while avoiding boundary misalignment due to structural variations, we implemented a scoring-based approach to identify conserved boundary Segments. The 40 Segments at the start and 40 Segments at the end of the pivot sub-Walk are used as candidate anchors. For each query assembly, a score is calculated for these candidate Segments based on: (i) their presence or absence in the query assembly, and (ii) the similarity between their step indices in the query compared to the pivot. The best-scoring segments define the conserved start and end boundaries for that query assembly. If no suitable boundaries are identified within the initial 40-segment windows, the search window shifts to the next 40 segments (forward from the start boundary, backward from the end boundary) until conserved anchors are found. Once boundaries are established, sub-walks are extracted for all query assemblies.

After sub-walk extraction, syntenic segments are identified through pairwise comparison (using node identifiers) between the pivot and each query assembly. Segments sharing the same identifier across assemblies are evaluated based on their Step positions (corrected by their respective starting Step). A distance score quantifies the Step shift between corresponding segments, accounting for local insertions or deletions that cause positional offsets. Segments are classified as syntenic when their distance score falls below a defined threshold (defined in the back-end), indicating conserved ordering despite minor rearrangements.

#### Annotation

Annotations can be loaded through the blue button consisting of six small squares located above the main view displaying the SVs, using either a **BED** (current pivot genome based coordinates) or a **GAF** (node based coordinates) file, preferably one generated by GrAnnoT [32]. Once loaded, clicking the same button allows to search by feature name, moving the view to the relevant locus.

When an annotation is loaded on the backend, we verify that the rows contain the expected columns; any that do not match are excluded. We then store everything in an in-memory dictionary on the server side. Searching for features by pattern takes *O*(1) time if the full feature name is specified or *O*(*n*) time — where *n* is the number of features — if a prefix or substring of the feature names is provided. For each query, the feature names matching the substring are potentially sorted twice; the cost of sorting N distinct names is *O*(*NlogN* ).

#### Diversity

The computation of the genetic divergence between individuals of a single or multiple PVG is performed using Mash [38]. As Mash requires FASTA sequence, we rebuild the FASTA sequences for all individuals by concatenating the Segments according to the different Walks. When completed, the FASTA sequences are submitted to *mash sketch* and *mash dist*. The resulting distance matrix is then analyzed by a multidimensional scaling approach (MDS) in order to reduce the dimension numbers to two and then visualized through an interactive scatterplot.

## Acknowledgments

The authors thank SELEOPRO, SYNGENTA, FLORIMOND DESPREZ, ARVALIS, MAS Seeds, GAUTIER SEMENCES SAS, RAGT 2n, and LIDEA for their financial support and their feedback during development, as well as INRAE and the PlantAlliance consortium for their logistical assistance within the framework of the SaVanache Project. The authors acknowledge also the ISO 9001 certified IRD i-Trop HPC at IRD Montpellier for providing HPC resources that have contributed to the research results reported within this paper. URL: https://bioinfo.ird.fr/.

## References

1. Richter BG, Sexton DP. Managing and Analyzing Next-Generation Sequence Data. PLOS Computational Biology. 2009 06;5(6):1–4. Available from: https://doi.org/10.1371/journal.pcbi.1000369. doi:10.1371/journal.pcbi.1000369.

2. Stephens ZD, Lee SY, Faghri F, Campbell RH, Zhai C, Efron MJ, et al. Big Data: Astronomical or Genomical? PLOS Biology. 2015 07;13(7):1–11. Available from: https://doi.org/10.1371/journal.pbio.1002195. doi:10.1371/journal.pbio.1002195.

3. Liu DX, Rajaby R, Wei LL, Zhang L, Yang ZQ, Yang QY, et al. Calling large indels in 1047 Arabidopsis with IndelEnsembler. Nucleic Acids Research. 2021 11;49(19):10879–94. Available from: https://doi.org/10.1093/nar/gkab904. arXiv:https://academic.oup.com/nar/article-pdf/49/19/10879/41070880/gkab904.pdf. doi:10.1093/nar/gkab904.

4. Shi L, Zhang P, Yu B, Cheng L, Liu S, Liu Q, et al. Genomic Analysis of Indel and SV Reveals Functional and Adaptive Signatures in Hubei Indigenous Cattle Breeds. Animals. 2025;15(12). Available from: https://www.mdpi.com/2076-2615/15/12/1755. doi:10.3390/ani15121755.

5. Schatz MC, Maron LG, Stein JC, Wences AH, Gurtowski J, Biggers E, et al. Whole genome de novo assemblies of three divergent strains of rice, Oryza sativa, document novel gene space of aus and indica. Genome Biology. 2014 Dec;15(11):506. Available from: https://doi.org/10.1186/s13059-014-0506-z. doi:10.1186/s13059-014-0506-z.

6. Golicz AA, Batley J, Edwards D. Towards plant pangenomics. Plant Biotechnology Journal. 2016;14(4):1099–105. Available from: https://onlinelibrary.wiley.com/doi/abs/10.1111/pbi.12499. arXiv:https://onlinelibrary.wiley.com/doi/pdf/10.1111/pbi.12499. doi:10.1111/pbi.12499.

7. Monat C, Sabot F. In: Rajora OP, editor. Pangenomics in Crop Plants. Cham: Springer International Publishing; 2024. p. 3–35. Available from: https://doi.org/10.1007/13836_2020_84. doi:10.1007/13836202084.

8. Nyaga DM, Zaied RE, Silander OK, Black MA, O’Sullivan JM. Beyond single references: pangenome graphs and the future of genomic medicine. Frontiers in Genetics. 2025;Volume 16 -2025. Available from: https://www.frontiersin.org/journals/genetics/articles/10.3389/fgene.2025.1679660. doi:10.3389/fgene.2025.1679660.

9. Medini D, Donati C, Tettelin H, Masignani V, Rappuoli R. The microbial pan-genome. Current opinion in genetics & development. 2005;15(6):589–94. doi:10.1016/j.gde.2005.09.006.

10. Garrison E, Guarracino A, Heumos S, Villani F, Bao Z, Tattini L, et al. Building pangenome graphs. Nature Methods. 2024 Nov;21(11):2008–12. Available from: https://doi.org/10.1038/s41592-024-02430-3. doi:10.1038/s41592-024-02430-3.

11. Secomandi S, Gallo GR, Rossi R, Rodríguez Fernandes C, Jarvis ED, Bonisoli-Alquati A, et al. Author Correction: Pangenome graphs and their applications in biodiversity genomics. Nature Genetics. 2025 Mar;57(3):763–3. Available from: https://doi.org/10.1038/s41588-025-02132-2. doi:10.1038/s41588-025-02132-2.

12. Bao Z, Weigel D. Complexity welcome: Pangenome graphs for comprehensive population genomics. Quantitative Plant Biology. 2025;6:e43. doi:10.1017/qpb.2025.10028.

13. Kopalli V, Arslan K, Morales-Díaz N, Zanini SF, Golicz AA. Toward a standardized framework for pangenome graph evaluation: assessing crop plant pangenome variation graph construction from multiple assemblies. GigaScience. 2025 01;14:giaf121. Available from: https://doi.org/10.1093/gigascience/giaf121. arXiv:https://academic.oup.com/gigascience/article-pdf/doi/10.1093/gigascience/giaf121/65738190/giaf121.pdf. doi:10.1093/gigascience/giaf121.

14. Hennig A, Bernhardt J, Nieselt K. Pan-Tetris: an interactive visualisation for Pan-genomes. BMC Bioinformatics. 2015 Aug;16(11):S3. Available from: https://doi.org/10.1186/1471-2105-16-S11-S3. doi:10.1186/1471-2105-16-S11-S3.

15. Pedersen TL, Nookaew I, Wayne Ussery D, Månsson M. PanViz: interactive visualization of the structure of functionally annotated pangenomes. Bioinformatics. 2017 04;33(7):1081–2. Available from: https://doi.org/10.1093/bioinformatics/btw761. arXiv:https://academic.oup.com/bioinformatics/article-pdf/33/7/1081/49038290/bioinformatics3371081.pdf. doi:10.1093/bioinformatics/btw761.

16. Ding W, Baumdicker F, Neher RA. panX: pan-genome analysis and exploration. Nucleic Acids Research. 2018 01;46(1):e5–5. Available from: https://doi.org/10.1093/nar/gkx977. arXiv:https://academic.oup.com/nar/article-pdf/46/1/e5/23327157/gkx977.pdf. doi:10.1093/nar/gkx977.

17. van den Brandt A, Jonkheer EM, van Workum DJM, van de Wetering H, Smit S, Vilanova A. PanVA: Pangenomic Variant Analysis. IEEE Transactions on Visualization and Computer Graphics. 2024;30(8):4895–909. doi:10.1109/TVCG.2023.3282364.

18. Wick RR, Schultz MB, Zobel J, Holt KE. Bandage: interactive visualization of de novo genome assemblies. Bioinformatics. 2015 10;31(20):3350–2. Available from: https://doi.org/10.1093/bioinformatics/btv383. arXiv:https://academic.oup.com/bioinformatics/article-pdf/31/20/3350/49036207/bioinformatics31203350.pdf. doi:10.1093/bioinformatics/btv383.

19. Gonnella G, Niehus N, Kurtz S. GfaViz: flexible and interactive visualization of GFA sequence graphs. Bioinformatics. 2019 08;35(16):2853–5. Available from: https://doi.org/10.1093/bioinformatics/bty1046. arXiv:https://academic.oup.com/bioinformatics/article-pdf/35/16/2853/50719178/bty1046.pdf. doi:10.1093/bioinformatics/bty1046.

20. Guarracino A, Heumos S, Nahnsen S, Prins P, Garrison E. ODGI: understanding pangenome graphs. Bioinformatics. 2022 07;38(13):3319–26. Available from: https://doi.org/10.1093/bioinformatics/btac308. arXiv:https://academic.oup.com/bioinformatics/article-pdf/38/13/3319/49883779/btac308.pdf. doi:10.1093/bioinformatics/btac308.

21. Beyer W, Novak AM, Hickey G, Chan J, Tan V, Paten B, et al. Sequence tube maps: making graph genomes intuitive to commuters. Bioinformatics. 2019 12;35(24):5318–20. Available from: https://doi.org/10.1093/bioinformatics/btz597. arXiv:https://academic.oup.com/bioinformatics/article-pdf/35/24/5318/48978177/bioinformatics35245318.pdf. doi:10.1093/bioinformatics/btz597.

22. Vorbrugg S, Bezrukov I, Bao Z, Weigel D. Gretl—variation GRaph Evaluation TooLkit. Bioinformatics. 2025 01;41(1):btae755. Available from: https://doi.org/10.1093/bioinformatics/btae755. arXiv:https://academic.oup.com/bioinformatics/article-pdf/41/1/btae755/61272645/btae755.pdf. doi:10.1093/bioinformatics/btae755.

23. Durant Sabot F, Conte M, Rouard M. Panache: a web browser-based viewer for linearized pangenomes. Bioinformatics. 2021 12;37(23):4556–8. Available from: https://doi.org/10.1093/bioinformatics/btab688. arXiv:https://academic.oup.com/bioinformatics/article-pdf/37/23/4556/50579116/btab688.pdf. doi:10.1093/bioinformatics/btab688.

24. Liu M, Zhang F, Lu H, Xue H, Dong X, Li Z, et al. PPanG: a precision pangenome browser enabling nucleotide-level analysis of genomic variations in individual genomes and their graph-based pangenome. BMC Genomics. 2024 Apr;25(1):405. Available from: https://doi.org/10.1186/s12864-024-10302-5. doi:10.1186/s12864-024-10302-5.

25. Computomics GmbH. Pantograph;. Available from: https://www.computomics.com/solutions/pantograph.html.

26. Miao Z, Yue JX. Interactive visualization and interpretation of pangenome graphs by linear reference–based coordinate projection and annotation integration. Genome Research. 2025. Available from: https://genome.cshlp.org/content/early/2025/02/05/gr.279461.124.abstract. doi:10.1101/gr.279461.124.

27. Jenike K, Brown N, Kovaka S, et al. Panagram: Interactive, Alignment-Free Pan-Genome Browser;. Available from: https://github.com/kjenike/panagram.

28. Mastromatteo S, Chirmade S, Roshandel D, Thiruvahindrapuram B, Wang Z, Patel RV, et al. PangyPlot: multi-scale interactive visualization of pangenome variation graphs. Bioinformatics. 2026 08;42(8):btag619. Available from: https://doi.org/10.1093/bioinformatics/btag619. arXiv:https://academic.oup.com/bioinformatics/article-pdf/42/8/btag619/70688669/btag619.pdf. doi:10.1093/bioinformatics/btag619.

29. Yokoyama TT, Sakamoto Y, Seki M, Suzuki Y, Kasahara M. MoMI-G: modular multi-scale integrated genome graph browser. BMC Bioinformatics. 2019 Nov;20(1):548. Available from: https://doi.org/10.1186/s12859-019-3145-2. doi:10.1186/s12859-019-3145-2.

30. Goel M, Schneeberger K. plotsr: visualizing structural similarities and rearrangements between multiple genomes. Bioinformatics. 2022 05;38(10):2922–6. Available from: https://doi.org/10.1093/bioinformatics/btac196. arXiv:https://academic.oup.com/bioinformatics/article-pdf/38/10/2922/49010748/btac196.pdf. doi:10.1093/bioinformatics/btac196.

31. Yokoyama TT, Kasahara M. Visualization tools for human structural variations identified by whole-genome sequencing. Journal of Human Genetics. 2020 Jan;65(1):49–60. Available from: https://doi.org/10.1038/s10038-019-0687-0. doi:10.1038/s10038-019-0687-0.

32. Marthe N, Zytnicki M, Sabot F. GrAnnoT, a tool for efficient and reliable annotation transfer through pangenome graph. Peer Community Journal. 2025;5. Available from: https://peercommunityjournal.org/articles/10.24072/pcjournal.651/. doi:10.24072/pcjournal.651.

33. Hickey G, Monlong J, Ebler J, Novak AM, Eizenga JM, Gao Y, et al. Pangenome graph construction from genome alignments with Minigraph-Cactus. Nature Biotechnology. 2024 Apr;42(4):663–73. Available from: https://doi.org/10.1038/s41587-023-01793-w. doi:10.1038/s41587-023-01793-w.

34. Bocs S, Carrette C, Confais J, Dubois S, Duvaux L, Klopp C, et al. Adopting graph-based pangenomics for agronomy and biodiversity studies: current resources and challenges. Peer Community Journal. 2026;6. Available from: https://peercommunityjournal.org/articles/10.24072/pcjournal.781/. doi:10.24072/pcjournal.781.

35. Harling-Lee JD, Gorzynski J, Yebra G, Angus T, Fitzgerald JR, Freeman TC. A graph-based approach for the visualisation and analysis of bacterial pangenomes. BMC Bioinformatics. 2022 Oct;23(1):416. Available from: https://doi.org/10.1186/s12859-022-04898-2. doi:10.1186/s12859-022-04898-2.

36. Dong X, Jiao D, Xue H, Fan S, Wei C. APAV: An advanced pangenome analysis and visualization toolkit. PLOS Computational Biology. 2025 07;21(7):1–10. Available from: https://doi.org/10.1371/journal.pcbi.1013288. doi:10.1371/journal.pcbi.1013288.

37. Durant E, Rouard M, Ganko EW, Muller C, Cleary AM, Farmer AD, et al. Ten simple rules for developing visualization tools in genomics. PLOS Computational Biology. 2022 11;18(11):1–11. Available from: https://doi.org/10.1371/journal.pcbi.1010622. doi:10.1371/journal.pcbi.1010622.

38. Ondov BD, Treangen TJ, Melsted P, Mallonee AB, Bergman NH, Koren S, et al. Mash: fast genome and metagenome distance estimation using MinHash. Genome Biology. 2016 Jun;17(1):132. Available from: https://doi.org/10.1186/s13059-016-0997-x. doi:10.1186/s13059-016-0997-x.

