## Supplementary material for "SaVanache: indexing and visualizing pangenome variation graphs": SuppTable 1

Table 1: Comparative table of pangenome visualization tools' representations and data

| Tool | Ref. | Vis. paradigms | Vis. abstraction & scale | Input data |
|---|---|---|---|---|
| Pan-Tetris | [1] | Gene set | Aggregable gene sets, colored by annotation | Ad-Hoc PanGee file + TIGRFAM annotations, PAV CSV |
| Bandage | [2] | Genome graph | Node-link sequence graph for small to medium-sized graphs, single or doubled nodes, optional BLAST hits overlaid | Assembly graph formats: LastGraph (Velvet), FASTG (SPAdes), Trinity.fasta, ASQG and GFA |
| PanViz | [3] | Gene set; Composite | PAV of genes, grouped by GO terms, in chord diagram, summary stacked bar charts and GO terms treemap | HTML built from PanVizGenerator (CSV or annotations from Blast2GO analysis) |
| GfaViz | [4] | Genome graph | Node-link sequence graph, configurable layouts, better suited for small to medium graphs | GFA files |
| panX | [5] | Gene set; Sequence alignment; Composite | Gene set summary dashboard (tables, trees, distributions...), MSAViewer for sequences | Built with `https://github.com/neherlab/pang-genome-analysis` from GenBank genome files |
| MoMI-G | [6] | Genome graph; Composite | Dashboard combining circular overview of SVs, list of SVs, and local SequenceTubeMap representation for nested SV inspection | vg-compatible format and annotation files (xg, GAM, CSV, BED) |
| SequenceTubeMap | [7] | Genome graph; Sequence alignment | Paths within a variation graph, local windows of nucleotide sequences, SNP, insertion and deletion marks on hover | GBZ variation graph (binary GFA subset), aligned reads in GAF/GAM |
| Panache | [8] | Anchored blocks; Browser | Navigation at 'panBlock' level, sequences abstracted out, chromosome-wide exploration | PAV CSV (on linear panreference coordinates), optional annotation GFF and Newick tree |
| ODGI | [9] | Plotter; Anchored blocks | Binned linear representation of graph paths and rearrangement structure of selected regions | Odgi graph, derived from GFA or vg toolkit output |
| plotsr | [10] | Plotter; Synteny | Consecutive pairwise comparisons of region/chromosome-wide synteny, color-coded SVs | Chromosome-level assemblies (FASTA) ran through pairwise structural annotations (from syri) and optional tracks (GFF, BED, bedGraph) |
| PPanG | [11] | Browser; Genome graph | Genome browser with additional gene annotation tracks and graph as a SequenceTubeMap view of a region, with exon mapping | Pangenome graph built with Cactus and pggb, annotated with MAKER |
| Pantograph | [12] | Anchored blocks; Browser; Composite | Karyotype view of QTLs; graph view with pre-computed bin widths | Pangenome graphs (format unclear), metadata (CSV), variations (VCF) |
| PanVA | [13] | Sequence alignment; Browser | Local navigation at nucleotide level per gene, phylogenetic tree, interactive grouping | Pangenome DB (PanTools): alignments, sequences, annotation... CSVs + optional tree |
| APAV | [14] | Gene set; Plotter; Composite | HTML reports with query/filter on PAV heatmap, overview summaries and complementary plots | Reference genome, GFF/BED regions, BAM alignments, optional sample metadata |
| VRPG | [15] | Genome graph; Composite | Coordinated views of local node-link graph around a backbone, gene and misc. annotations | rGFA, GFA, annotation tracks in GFF and BED formats |
| PanGraphViewer | [16] | Genome graph | Node-link diagram of local subgraph, variations from backbone genome encoded through node shape | graph rGFA, GFA, variation VCF, annotation BED or GFF |
| PangyPlot | [17] | Genome graph; Browser; Composite | Coordinated views of chromosome diversity, locus selector and local pangenome graph with highlighted annotations | GFA graph, prepped through vg/odgi |
| Panagram | [18] | Plotter; Browser; Composite | Pangenome summary overview to chromosome inspection of pan-k-mer conservation bitmaps | TSVs of genome FASTA and optional GFF annotation for k-mer construction |

Table 1: Comparative table of pangenome visualization tools' representations and data

| Tool | Ref. | Vis. paradigms | Vis. abstraction & scale | Input data |
|---|---|---|---|---|
| SaVanache | - | Anchored blocks; Browser; Composite | Pangenome diversity overview, SV browser using glyphs for one-to-many comparisons against selected pivot, down to local alignment views | 'minigraph-cactus'-like GFA, BED or GAF annotations (preferably from GrAnnoT) |
