## Supplementary figures and images for "SaVanache: indexing and visualizing pangenome variation graphs"

### S Figure 1.png

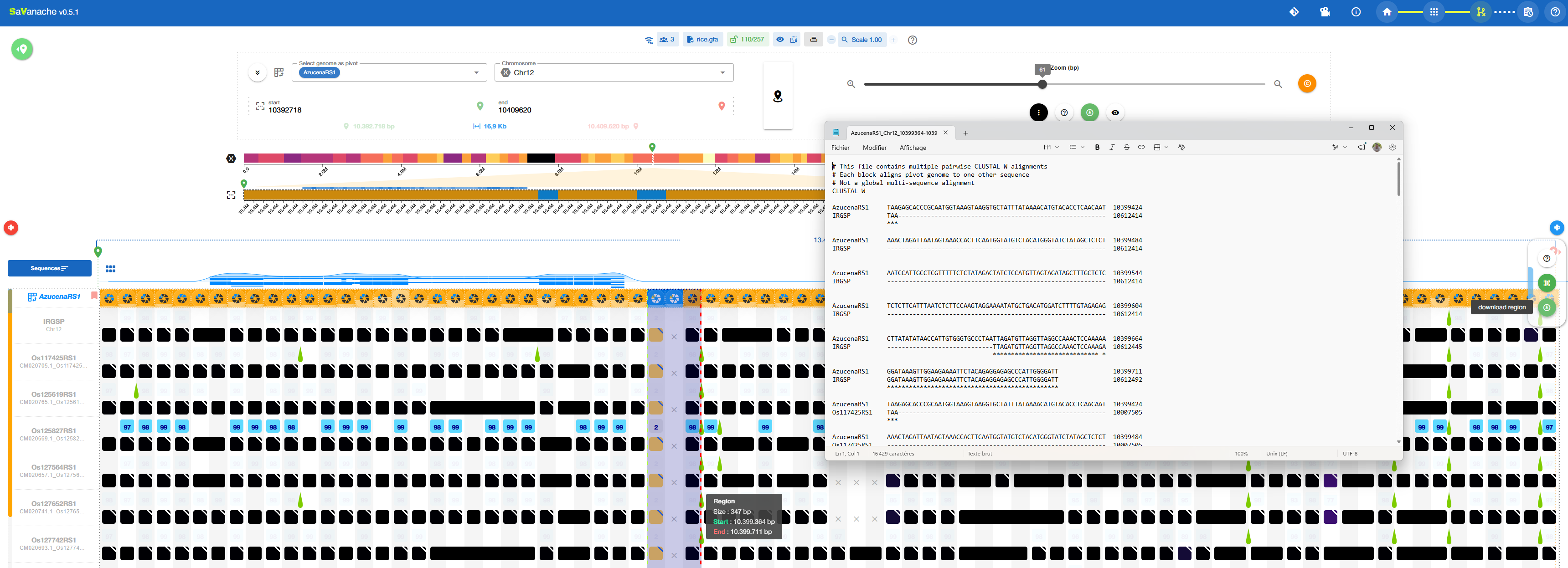
